# A NET4-RabG3 couple mediate the link between actin and the tonoplast and is essential for normal actin cytoskeletal remodelling in stomatal closure to flg22

**DOI:** 10.1101/2021.09.29.461190

**Authors:** Timothy J. Hawkins, Michaela Kopischke, David A. Mentlak, Patrick Duckney, Johan T.M. Kroon, Mai Thu Bui, A. Christine Richardson, Mary Casey, Agnieszka Alexander, Geert De Jaeger, Monika Kalde, Ian Moore, Yasin Dagdas, Patrick J. Hussey, Silke Robatzek

**Affiliations:** Department of Biosciences, University of Durham, South Road, Durham, DH1 3LE, UK; The Sainsbury Laboratory, Norwich Research Park, NR4 7UH, UK; LMU Munich Biocenter, Großhadener Strasse 4, 82152 Planegg, DE; Gregor Mendel Institute (GMI), Austrian Academy of Sciences, Vienna BioCenter, Vienna, AUT; VIB-University Ghent, Center for Plant System Biology, Technologiepark 927, 9052 Ghent, BE; Department of Plant Sciences, University of Oxford, South Parks Rd., Oxford, OX1 3RB, UK

**Keywords:** NET4, RABG3, flg22, PAMP, actin, vacuole, tonoplast, stomata

## Abstract

Members of the NETWORKED (NET) family are involved in actin-membrane interactions. They tether the cell’s plasma membrane (PM) to the actin network. Moreover, in a similar manner, they are also involved in the tethering of membrane bound organelles to the actin cytoskeleton; the endoplasmic reticulum (ER) and the ER to the PM. This raises the question as to whether NET proteins are involved in actin cytoskeletal remodelling. Here we show that two members of the NET family, NET4A and NET4B, are essential for normal guard cell actin reorganization, which is a process critical for stomatal closure in plant immunity. NET4 proteins interact with F-actin and with members of the Rab7 GTPase RABG3 family through two distinct domains, allowing for simultaneous localization to actin filaments and the tonoplast. NET4 proteins interact with GTP-bound, active RABG3 members, suggesting their function as downstream effectors. We also show that RABG3b is critical for stomatal closure induced by microbial patterns. Taken together, we conclude that the actin cytoskeletal remodelling during stomatal closure depends on a molecular link between actin filaments and the tonoplast, which is mediated by the NET4-RABG3b interaction. We propose that stomatal closure to microbial patterns involves the coordinated action of immune signalling events and proper actin cytoskeletal remodelling.

## Introduction

The NETWORKED (NET) family of proteins are key molecular components of actin-membrane interactions in plants ^1, 2^. As such, they directly bind to the actin cytoskeleton and localise to membrane compartments. Different NET proteins link actin to specific membranes, including actin-vacuole interactions (NET4A), actin-nuclear membrane interactions (NET3A), actin-plasma membrane (PM) interactions (NET2A), actin-plasmodesmata interactions (NET1A), actin-endoplasmic reticulum (ER) interactions (NET3B), and actin-PM/ER contact site (EPCS) interactions (NET3C) ^1, 3, 4^. These actin-membrane interactions are thought to influence actin dynamics and membrane morphology, responsible for controlling root and pollen tube development, cell expansion and pavement cell morphogenesis ^1, 5, 6, 7^.

Underpinning these processes are interactions of NET proteins with other proteins to form complexes that could be tissue or developmental stage specific, overlapping with the expression patterns of the *NET* genes. For example, NET2A is expressed specifically in pollen and forms a complex with the transmembrane POLLEN RECEPTOR-LIKE KINASES (PRK) 4 and 5, indicating that NET proteins could link external signalling events with actin-membrane dynamics ^5^. NET3C exists in complexes with VAP27, KINESIN-LIGHT-CHAIN-RELATED (KLCR) and calmodulin-binding IQ67 DOMAIN (IQD) proteins, which mediate the interaction between actin, microtubules and ER at the EPCS ^8, 7^. It is thus possible that EPCS represent focal points of signal sensing and cytoskeletal organization. Given their link to the cytoskeleton and specific lipid and protein composition, EPCS could also have roles in membrane trafficking, i.e. endocytosis and/or autophagy ^2, 9^.

Both NET1A and NET4A are predominantly expressed in the root meristem and elongation zone ^1^. However, NET1A and NET4A localize to distinct membranes, the plasmodesmata and the vacuole respectively, and therefore they likely control actin-membrane dynamics in different plant processes. NET1A is involved in root development but the process regulated by NET4A remains unknown ^1^. Given that NET4A localizes to the vacuole, it is interesting to note that cells of the root meristem and elongation zone are characterized by complex vacuolar structures, i.e. fragmented and/or with tubular morphology ^10^. Only recently it was reported in roots that NET4A appears to have a stronger vacuolar localization in the centre of the cell, a region of higher vacuolar constrictions. Knock-out mutants of NET4A and its closest homologue NET4B, and NET4A overexpression result in more spherical vacuoles ^11^.

Morphological dynamics of vacuoles are not only involved in plant growth but are a hallmark of stomatal movements ^12^. The vacuoles in the two guard cells that form the stomatal pore change their volume and size, driving the closure and opening of the pore. When stomata close, the vacuole morphology becomes more complex with invaginations and bulb-like structures in the lumen, increasing the total vacuole surface area by 20% ^12^. These intra-vacuolar structures could represent a storage of excess membrane material during stomatal closure.

Actin filament dynamics are associated with stomatal movements, forming distinct actin arrays within guard cells: radial orientation in open stomata and a longitudinal orientation in closed stomata ^13^. Actin filaments transiently bundle during stomatal opening but bundles dissolve when stomata are fully open. Consequently, chemical interference with actin dynamics affects stomatal movements. Blocking actin polymerization with Cytochalasin D promotes light-induced stomatal opening, while Latrunculin B promotes abscisic acid (ABA)-induced stomatal closure ^14, 15^. Stabilizing actin with Phalloidin and Jasplakinolide inhibits ABA-induced stomatal closing ^15^. Genetic disruption of actin bundling, i.e. upon overexpression of the ACTIN DEPOLYMERIZATION FACTOR 1 (ADF1) or inactivation of the small GTPase AtRac1 induces stomatal closure ^16, 17^.

Guard cells respond to numerous physiological and environmental cues, including stomatal closure induced upon drought stress and pathogen attack ^18^. Thus, signalling events are likely impacting on guard cell actin dynamics. Perception of pathogen-associated molecular patterns (PAMPs) such as bacterial flagellin and fungal chitin closes stomata and thereby protect against pathogen invasion ^18^. The cognate receptor for flagellin and its derived peptide flg22 is FLAGELLIN SENSING 2 (FLS2), which is highly expressed in guard cells ^19, 20^. Ligand-activation of the FLS2 – BRASSINOSTEROID INSENSITIVE 1-ASSOCIATED RECEPTOR KINASE 1 (BAK1) receptor complex induces signalling cascades resulting in the phosphorylation of the RESPIRATORY BURST OXIDASE HOMOLOG PROTEIN D (RBOHD) NADPH oxidase, S-type anion channel SLOW ANION CHANNEL1 (SLAC1) and its homologue SLAH3, required for PAMP-triggered stomatal closure ^21, 22^.

Additionally, flg22 perception results in a rapid change in the higher ordered state of the actin filament array within the guard cell, mostly showing radial array and radial bundles during the treatment ^13^. Guard cell actin filament orientation and density remained unchanged upon flg22-induced stomatal closure. However, epidermal pavement cells show increased actin filament abundance in response to flg22 and chitin in a FLS2, CHITIN ELICITOR RECEPTOR KINASE1 (CERK1) and ADF4-dependent manner ^23, 24^, suggesting cell type-specific differences in PAMP-induced actin filament dynamics. Whether actin-membrane interaction, such as NET4A-mediated actin-vacuole tethering, play roles in PAMP-induced immune responses, including stomatal closure, remains to be determined.

In this work, we examined NET4A and its closest homologue NET4B. They both localized to actin filaments and the tonoplast. Expression studies and phenotypic analysis of knock-out mutants revealed a role for both of the NET4 proteins in stomatal closure to flg22, indicative of their similar function. The *net4* mutants exhibit impaired stomatal closure at long treatment times but at earlier time points they were not immune-compromised. Using a combination of yeast-two-hybrid (Y2H), co-immunoprecipitation and FRET-FLIM assays, we show that the NET4 proteins form homo-and heterodimeric complexes, and interact with members of the RABG3 family. We also demonstrate that *rabg3b* mutants, but not mutants in other *RABG3* genes, are specifically impaired in flg22-induced stomatal closure, phenocopying the two *net4* mutants. In addition, analysis of the subcellular function of NET4 and RABG3 using the *net4* and *rabg3b* mutants suggest that the couple is involved in the reorganisation of the actin cytoskeleton. Taken together, these data support a model of the two NET4 proteins as RABG3B effectors, regulating guard cell actin cytoskeletal remodelling in response to PAMP-triggered stomatal closure.

## Results

### NET4 proteins localize to actin filaments and the tonoplast

Previous work suggested NET4A localizing to the actin cytoskeleton and the vacuole ^1, 11, 25^. We further defined the subcellular localization of NET4A. Confocal microscopy of stable transgenic *Arabidopsis thaliana* lines expressing NET4A-GFP under its native promoter exhibited detectable signals in roots and revealed a striking filamentous network pattern of NET4A-GFP (Fig. 1A). Co-expression with the actin marker Lifeact-RFP confirmed the localization of NET4A-GFP to the actin cytoskeleton (Fig. 1A). Detailed inspection showed discrete NET4A-GFP punctae labelling along the Lifeact-RFP actin filaments (Fig. 1A; Movie S1), a characteristic “beads-on-a-string” localization similar to other members of the NET family ^1^. Furthermore, analysis of transgenic Arabidopsis lines co-expressing native promotor-driven NET4A-GFP and the well-established tonoplast marker RABG3f-mCherry showed that NET4A localizes at the tonoplast (Fig. 1B 1C; Movies S2, S3) ^26^.

**Figure 1.**
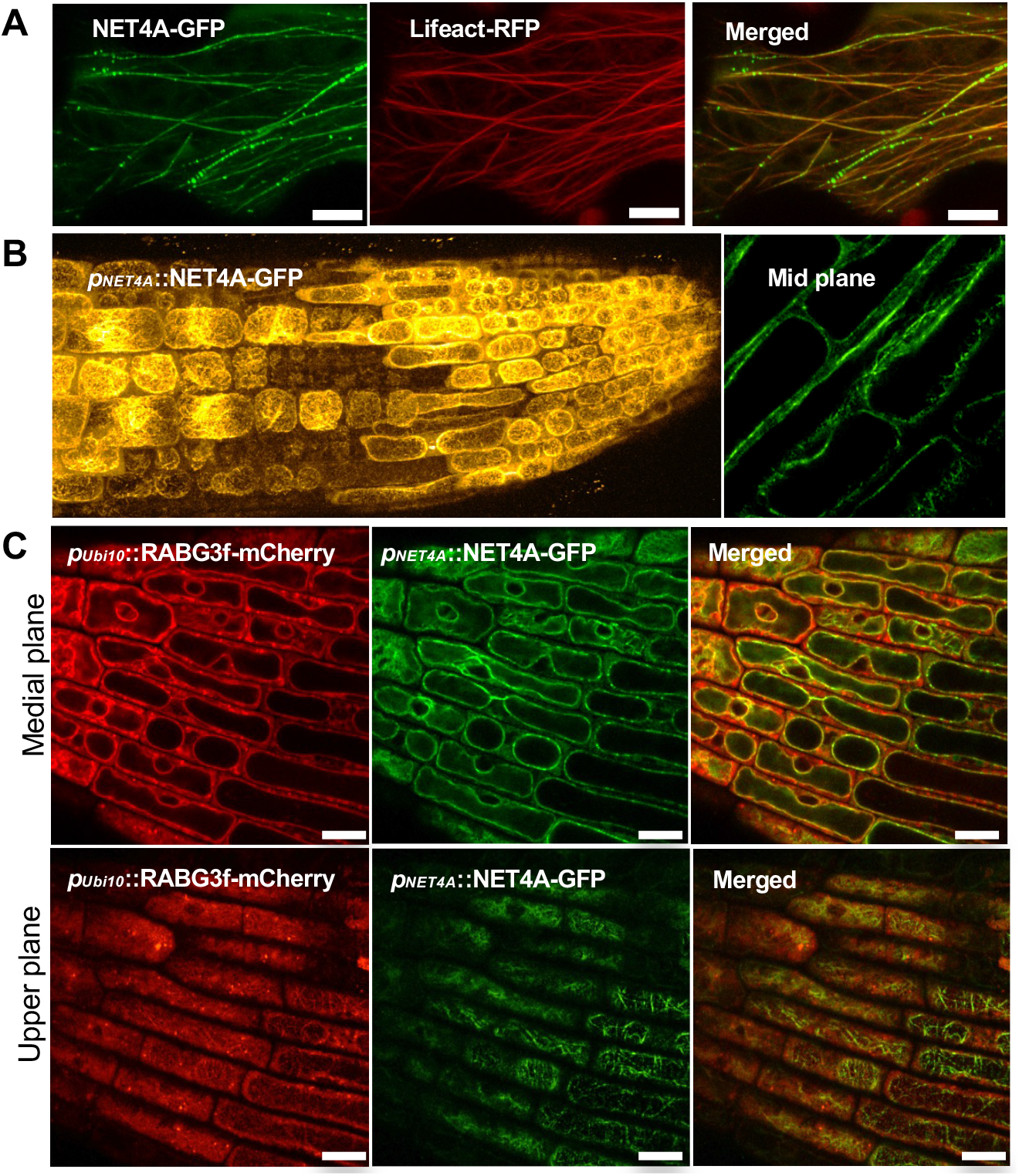
NET4 proteins decorate actin filaments and localise to the tonoplast. **(A)** Confocal microscopy of *N. benthamiana* leaves transiently expressing NET4A-GFP and Lifeact-RFP; scale bar = 10 μm. **(B)** Confocal microscopy of transgenic Arabidopsis lines expressing NET4A-GFP under its native promoter. NET4A-GFP localises to filaments and the tonoplast membrane; scale bar = 20 μm. **(C)** Confocal microscopy of transgenic Arabidopsis lines co-expressing *pNET4A*::NET4A-GFP and pUbi10::mCherry-RABG3f. NET4A-GFP and mCherry-RABG3f co-localise at the tonoplast *in planta*; scale bars = 10 μm.

Since NET4B-GFP expression was below the detection limit in stable transgenic Arabidopsis lines, we studied its localization in *Nicotinana benthamiana*. Transient expression of NET4B-GFP showed patterns reminiscent of NET4A-GFP and co-localization with the actin marker FABD2-mCherry (S1A). A hallmark of NET family proteins is the NET actin-binding (NAB) domain ^1^. To corroborate that the observed association of NET4B with actin represents direct binding to microfilaments, we mixed the recombinant NET4B 6xHIS-NAB domain with purified actin in a co-sedimentation assay. Upon ultracentrifugation, 6HIS-NET4B^1-105^ co-sedimented in the presence of F-actin, proving a direct association between NET4B and microfilaments (Fig. S1B). Using an isotype-specific anti-NET4B antibody and immuno-gold labelling, we detected enriched signals in close proximity to tonoplast structures of lytic vacuoles by transmission electron microscopy (Fig. S1C). Taken together, we conclude that NET4 proteins directly bind actin through the NAB domain and at the same time associate with the tonoplast. This simultaneous localization pattern is consistent with the role of NET family proteins as actin-membrane tethers.

Being close homologues and exhibiting the same localization patterns, we hypothesized that NET4A and NET4B could co-localize and interact with each other. Concordantly, confocal microscopy of NET4A-GFP and NET4B-RFP in *N. benthamiana* revealed their co-localization, observed as actin cytoskeleton patterns (Fig. 2A). Consistently, we found that both NET4A and NET4B can form homo- and heterodimers in yeast (Fig. 2B), and *in planta* (Fig. 2C). FRET-FLIM analysis of co-expressing NET4A-GFP and NET4B-RFP *N. benthamiana* leaves confirmed this interaction through the significant reduction in donor lifetime (Fig. 2D, 2E).

**Figure 2.**
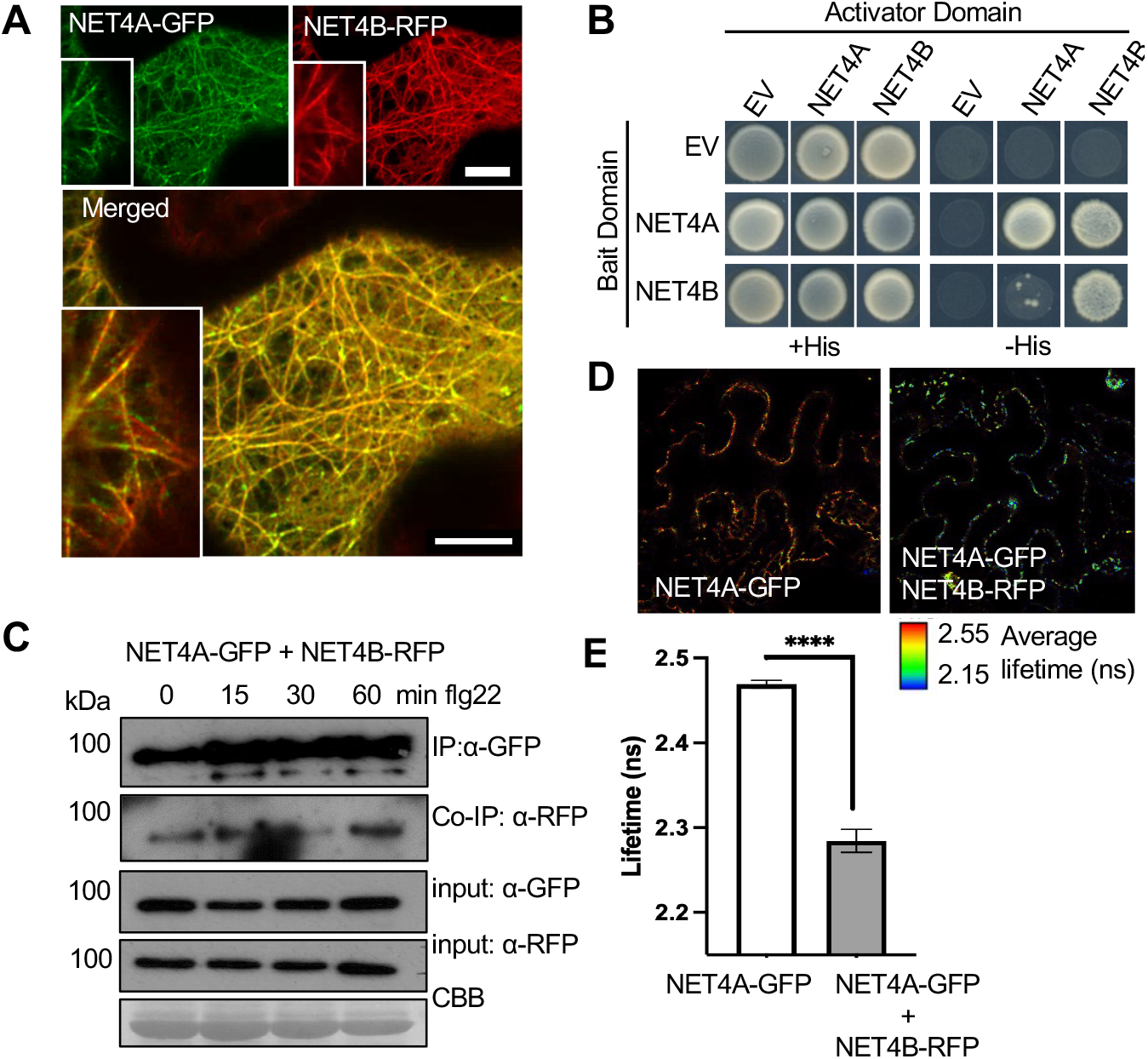
NET4 proteins co-localize, interact with each other and themselves. **(A)** Confocal micrographs of *N. benthamiana* leaf epidermal cells transiently co-expressing NET4A-GFP and NET4B-RFP. Shown are single plane confocal images capturing GFP fluorescence (green), RFP fluorescence (red) and overlaid signals (yellow); scale bars = 20 μm. **(B)** Yeast-two-hybrid analysis of the indicated constructs indicate homo- and heterodimer interactions of NET4A and NET4B (EV = empty vector). **(C)** Co-immunoprecipitation (Co-IP) assay of NET4A-GFP with NET4B-RFP. Fusion proteins were co-expressed in *N. benthamiana*, immunoprecipitated (IP) using GFP-trap beads and detected using fluorophore fusion specific antibodies (GFP and RFP). 1% of the input is shown as loading control. Blots are representative of 2 independent experiments. **(D)** FRET-FLIM analysis of NET4A-GFP and NET4B-RFP interaction. FLIM (TCSPC) confocal images of *N. benthamiana* leaf epidermal cells co-expressing both NET4 proteins. Data shown using lifetime LUT. **(E)** Graph of fluorescence lifetime values of donor alone and donor + acceptor. The significant reduction in lifetime of the donor in the presence of the acceptor indicates energy transfer and a physical interaction; n = 10 p <0.0001 Mann Whitney t-test.

### NET4 proteins are required for proper stomatal closure to flg22

To gain insights into the plant processes, which could be regulated by NET4A and NETB, we evaluated their transcriptional expression patterns using stable promoter GUS lines in *A. thaliana*. As shown in Fig. 3, we observed clear GUS signals in roots across cell files and developmental zones for both promoters (Fig. 3A, 3B). This is in agreement with the previously reported expression of NET4A-GFP in root epidermal cells ^1, 27^ 11. In addition, we found GUS expression in trichomes conferred by the *NET4A* promoter and in guard cells by the *NET4B* promoter. To validate *NET4* guard cell expression, we performed qRT-PCR analysis and found a high expression of *NET4B* but not *NET4A* in guard cell enriched samples compared to mesophyll cells (Fig. 3C). Analysis of AtGenExpress and GEO microarray experiment data confirmed *NET4B* expression in guard cells and revealed transcriptional responsiveness to flg22 and bacterial infection (*NET4A* is not available on the Affymetrix Arabidopsis ATH1 genome array used in these experiments).

**Figure 3.**
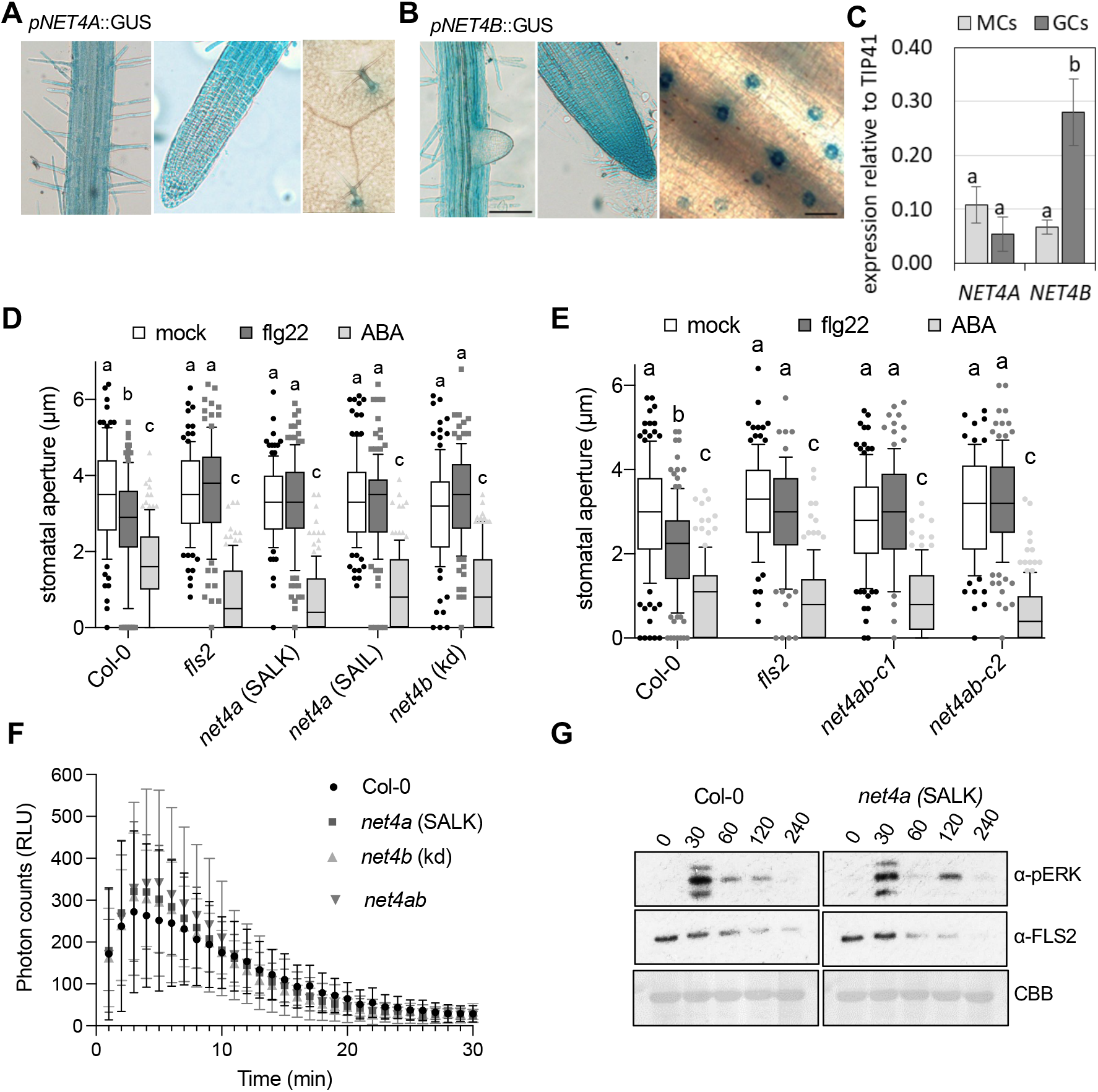
*net4* mutants are impaired in flg22- but not ABA-induced stomatal closure and other PTI responses. **(A, B)** GUS-stained tissues of stable transgenic Arabidopsis *NET4A* and *NET4B* promoter-driven GUS reporter lines. **(C)** Transcript levels of *NET4A* and *NET4B* were quantified in guard cells (GC)-enriched samples or mesophyll cells (MC), respectively, relative to *TIP41*. Bars represent mean values ± SEM of 3 biological replicates (ANOVA with Bonferroni’s Multiple Comparison Test). Different letters indicate significantly different values at p < 0.05. **(D, E)** Stomatal aperture measurements in *net4a* and *net4b* single and *net4ab-c1 and -c2* CRISPR-Cas double mutants compared with Col-0 and *fls2*. Stomatal apertures measured 2 hr after treatment with 20 μM flg22 and 10 μM ABA. Box plots of the values are shown with whiskers from the 5th to 95th percentiles, the line in the box shows the median (**(A)** n = 93-118 stomata (mock), n = 102-126 stomata (flg22), n = 113-125 stomata (ABA); **(B)** n = 113-131 stomata (mock), n = 92-124 stomata (flg22), n = 109-133 stomata (ABA)). Different letters indicate significantly different values at p < 0.0001 (2-way ANOVA, multiple comparisons). Both experiments were repeated twice with similar results. **(F)** ROS production measured as relative luminescence units (RLU) in the indicated genotypes treated with 100 nM flg22 over time. Graph represents ±SEM; n = 12 leaf discs. Experiment is representative for 2 independent experiments with similar results. **(G)** Flg22-induced activation of MAPK in the indicated genotypes and time points. MAPK activation is revealed with anti-pERK antibodies. For control, FLS2 abundance and Coomassie brilliant blue (CBB) staining is shown. The experiment was repeated twice with similar results.

Next, we isolated T-DNA insertional lines for both *NET4* genes resulting in loss-of-function of *NET4A* and knock-down of *NET4B*, selected *net4a* x *net4b* double mutants from single line crosses, and generated two *net4ab* double null mutants using CRISPR-Cas, all used for phenotype characterization (Fig. S2A, S2B). Although previous works point at roles for NET4A and NET4B in vacuoles morphology in roots ^11^, we did not observe obvious defects in plant growth and development (Fig. S2C). Because previous work identified that stomatal guard cell actin filament organization is altered in response to flg22 ^23, 28^, we focussed our phenotype analysis on stomatal behaviours. Single and double *net4a* and *net4b* mutants exhibited impaired stomatal closure to flg22 but not ABA (Fig. 3D, 3E), suggesting a specific impairment in pattern-triggered immunity (PTI). However, *net4* mutants were not generally immune-compromised, showing wild type-like production of reactive oxygen species (ROS) and activation of mitogen-activated protein kinases (MAPKs) (Fig. 3F, 3G), two typical PTI responses and ROS being required for stomatal closure ^18, 21^. NET4A and NET4B homo- and heterodimer formation was independent of flg22 (Fig. 2C). Thus, NET4A and NET4B do not appear to be required for PTI signalling but rather could be involved in cellular aspects of stomatal closure.

### NET4 proteins behave like downstream effectors of RABG3 GTPases

To further identify functions of NET4A and NET4B, we screened for interacting proteins using tandem-affinity purification followed by mass-spectrometry analysis (Table S1). In yeast, both, NET4A and NET4B interacted with members of the RABG3 family, including RABG3a and RABG3f (Fig. 4A). Multiple members of the RABG3 class localise to the tonoplast and have roles in vacuolar trafficking and fusion ^29, 26^. Therefore, we focused on RABG3 family proteins as interactors of NET4A and NET4B. We validated their interaction *in planta*, detecting association of NET4B with YFP-RABG3f in immunoprecipitates analysed by mass-spectrometry (Table S2). The protein sequences of RABG3 members are very similar, which may account for the ability of NET4 proteins to interact with multiple isoforms. The relatively large number of branches within the RabG class in Arabidopsis may reflect the diversity of vacuoles and associated trafficking that have evolved in higher plants ^25^.

**Figure 4.**
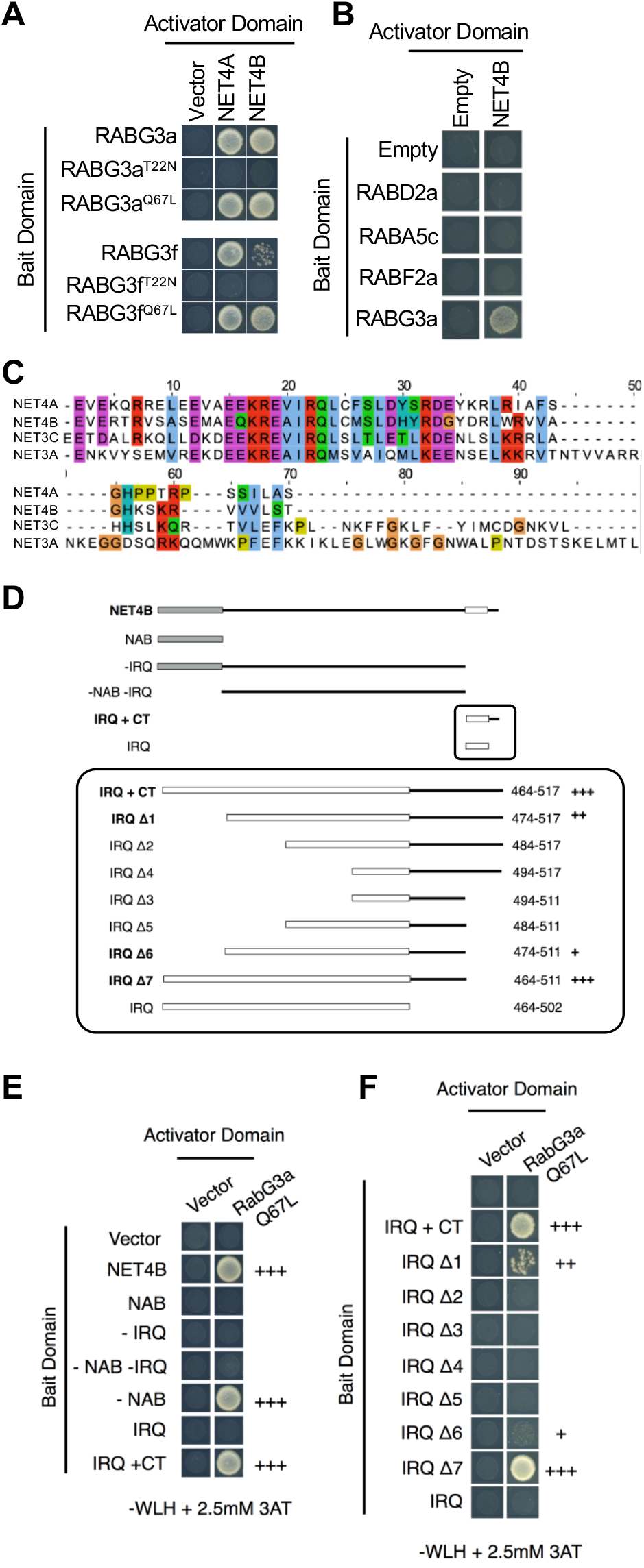
NET4 proteins interact with RabG3 GTPases in an activation- and motif-dependent manner. **(A)** Yeast-two-hybrid analysis of NET4A and NET4B indicate interactions with constitutive active (CA = Q67L) but not dominant-negative (DN = T22N) RABG3 members. **(B)** Yeast-two-hybrid analysis of NET4A and NET4B shows no interactions with members of other Rab GTPase families than RABG3. **(C)** Sequence alignment of IRQ domains including the C-terminus (CT). Of all NET family members, IRQ domains are only found in NET4A, NET4B, NET3A and NET3C. **(D)** Overview of truncated NET4B proteins used to identify the minimal interaction domain required for interaction with RABG3 proteins. **(E, F)** Yeast-two-hybrid analysis of the indicated truncated NET4B proteins with constitutive active (CA = Q67L) RABG3a; NAB = NET actin binding domain; IRQ = conserved sequence motif; CT = C-terminus.

RABG3 family proteins are Rab7 GTPases, which cycle in an active, GTP-bound state and an inactive, GDP-bound state, which are determinants of their subcellular localization and likely interaction partners ^30^. We therefore tested the interaction of NET4A and NET4B with both constitutive active (CA) and GDP-locked (TN) dominant negative (DN) variants of RABG3a and RABG3f. Only wild type and active, GTP-bound RABG3a and RABG3f interacted with NET4A and NET4B (Fig. 4A). We then addressed whether the association of NET4 proteins is specific to the RABG proteins or whether this interaction occurs more broadly across the Arabidopsis Rab GTPase families. Using RABD2a, RABA5c and RABF2a as baits, we identified no interaction with NET4B in Y2H (Fig. 4B), and similarly NET4B was not co-immunoprecipitated from Arabidopsis microsomes expressing YFP-RABA2a or YFP-RABA5c (Table S2). Since activation-dependent interactions, in a GTP-bound manner, are characteristic for Rab GTPase effectors ^30^, we suggest that NET4 proteins represent downstream effectors of RABG3 GTPases.

### NET4 proteins contain a novel RABG3 specific binding motif

The C-terminus of NET4B contains a conserved domain with an invariable “IRQ” sequence, thus referred to as the IRQ domain (Fig. 4C). C-terminal deletion including the IRQ domain abolished the interaction of NET4B with CA-RABG3a (Fig. 4D, 4E). To define the minimal domain of the NET4B C-terminus that is required for the interaction with RABG3a, we tested a series of truncated IRQ domains with and without the C-terminus as Y2H baits against CA-RABG3a (Fig. 4D). This experiment identified residues 464-511 (47aa) of NET4B as the RABG3 binding domain, a novel RABG3-specific binding motif previously unknown in plants (Fig. 4F). We suggest that NET4 proteins associate to the tonoplast, because of their interaction with GTP-bound RABG3 GTPases through the IRQ domain. In agreement, we found that the interaction between RABG3a and NET4B was independent of its NAB domain, using a NET4B NAB deletion variant and the NAB domain alone as Y2H baits with CA-RABG3a as the prey (Fig. 4D, 4E).

### RABG3b is required for proper flg22-induced stomatal closure

Having observed that NET4A and NET4B mediate stomatal closure to flg22, we determined the guard cell expression levels of all RABG3 family proteins by qRT-PCR. Although RABG3B, RABG3F and RAGB3E show high guard cell expression, the levels were not higher than those in mesophyll cells (Fig. 5A). Next, we obtained T-DNA insertional lines for all *RABG3* genes and screened the mutants for stomatal responses. Consistent with previous findings ^31^, stomata from *rabg3b* mutants did not close in response to flg22 (Fig. 5B). This phenotype of stomatal closure was specific to RABG3b as none of the other tested *rabg3* mutants, including a *rab2g* mutant, showed altered closure of stomata to both flg22 and ABA (Fig. 5B). Phylogenetic analysis of the RabG3 subclass places the isoforms into two major branches, one containing RabG3c, RABG3d, RabG3e and RabG3f and a second contain RabG3a and b ^32, 33^. This structural distinction may in part explain why of those RABG proteins highly expressed in guard cells only *rabg3b* exhibits altered stomatal closure.

**Figure 5.**
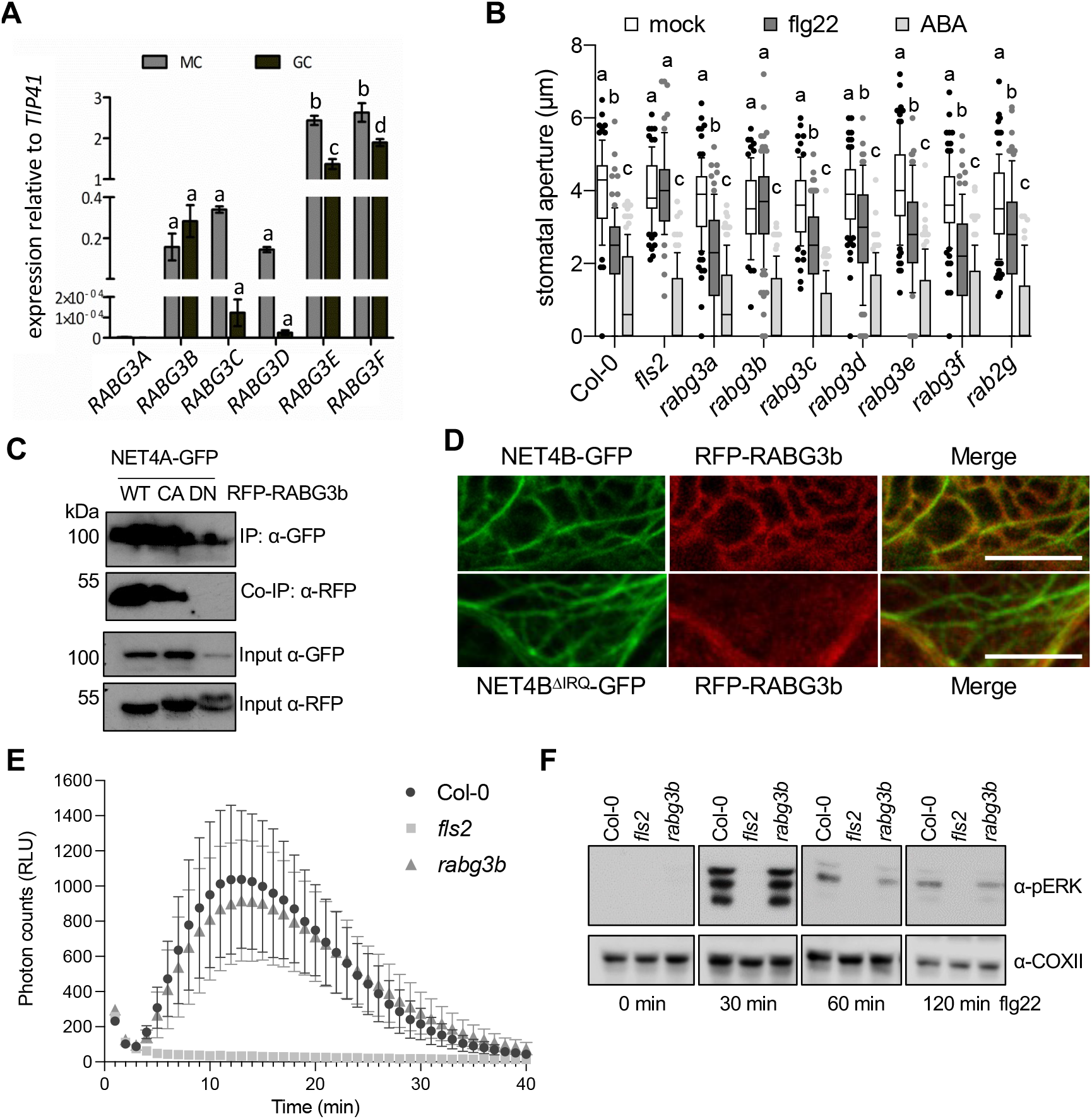
*rabg3b* mutants are impaired in flg22- but not ABA-induced stomatal closure. **(A)** Transcript levels of *RABG3* family members were quantified in guard cells (GC)-enriched samples or mesophyll cells (MC), respectively, relative to *TIP41*. Bars represent mean values ± SEM of 3 biological replicates (ANOVA with Bonferroni’s Multiple Comparison Test). **(B)** Stomatal aperture measurements in *rabg3* and *rabg2* mutants, Col-0 and *fls2*. Apertures measured 2 hr after treatment with 20 μM flg22 or 10 μM ABA. Box plots of the values are shown with whiskers from the 5th to 95th percentiles, the line in the box shows the median (n = 77-105 stomata (mock), n = 69-105 stomata (flg22), n = 105-117 stomata (ABA)). Different letters indicate significantly different values at p < 0.0001 (2-way ANOVA, multiple comparisons). The experiment was repeated twice with similar results. **(C)** Co-immunoprecipitation (Co-IP) assay of NET4A-GFP with wild type (WT), constitutive active (CA) and dominant-negative (DN) RFP-RABG3b. 1% of the input is shown as loading control. **(D)** Confocal microscopy of *N. benthamiana* leaves transiently expressing RFP-RABG3b and NET4B-GFP or NET4B^ΔIRQ^-GFP; scale bar = 10 μm. **(E)** ROS production measured as relative luminescence units (RLU) in the indicated genotypes treated with 100 nM flg22 over time. Graph represents ±SEM; n = 3 leaf discs. The experiment was repeated twice with similar results **(F)** Flg22-induced activation of MAPK in the indicated genotypes and time points. MAPK activation is revealed with anti-pERK antibodies. For loading control, COXII abundance is shown. The experiment was repeated twice with similar results.

To validate our observations, we developed complementation lines. Immunoblot analysis of constitutively expressed GFP-RABG3b in the *rabg3b* background revealed signals of full-length GFP-RABG3b fusion proteins and free GFP (Fig. S3A). However, confocal microscopy revealed strong signals at tonoplast structures in pavement and guard cells (Fig. S3B). This is consistent with the previously described localization of other RABG3 family members ^26^, including RABG3f (Fig. 1C), and suggests that the accumulation of free GFP in the immunoblot analysis could be due to the tissue disruption. Of relevance, GFP-RABG3b expression was functionally complementing the impaired flg22-induced stomatal closure in the *rabg3b* mutant (Fig. S3C).

To investigate whether RABG3b associates with NET4 proteins, we performed co-immunoprecipitation analysis. Co-expression of NET4A-GFP and RFP-RABG3b in *N. benthamiana* showed an association of NET4A with wild type RABG3b as well as with CA-RABG3b but not DN-RABG3b (Fig. 5C). Additional Y2H screening using CA-RABG3b as a bait identified NET4A as most prominent interactor (Table S3). This is consistent with our results on NET4 protein interaction with RABG3a and RABG3f (Fig. 4A), and further highlights that NET4 proteins represent effectors of RABG3 family members. Co-expression of NET4B-GFP and RFP-RABG3b in *N. benthamiana* showed signals of RABG3b-labelled membrane structures at NET4B-labelled filaments. This co-localization was not observed during co-expression of NET4B IRQ-GFP which lacks the IRQ domain, demonstrating *in planta* that the IRQ domain is required for the association of NET4B and RABG3 (Fig. 5D), in agreement with our Y2H findings (Fig. 4F).

Phenocopying the *net4* mutants, *rabg3b* was not generally immune-compromised, exhibiting wild type-like flg22-induced ROS production and MAPK activation (Fig. 5E, 5F), and did not show obvious defects on plant growth and development (Fig. S3D). This is consistent with previous findings, reporting the requirement of *rabg3* higher order mutants to observe phenotypes ^26^. Stomatal closure represents a PTI response ^18^. Thus, the impaired stomatal closure of *net4* and *rabg3b* mutants contrasts their wild type-like ROS and MAPK induction to flg22. We therefore evaluated the stomatal response to flg22 at two different time points. We detected that the *net4* and *rabg3b* mutants showed wild type-like stomatal closure after 1 hr flg22 treatment but after 2 hrs, mutant stomata were open, while wild type stomata were closed (Fig. 6A, 6B).

**Figure 6.**
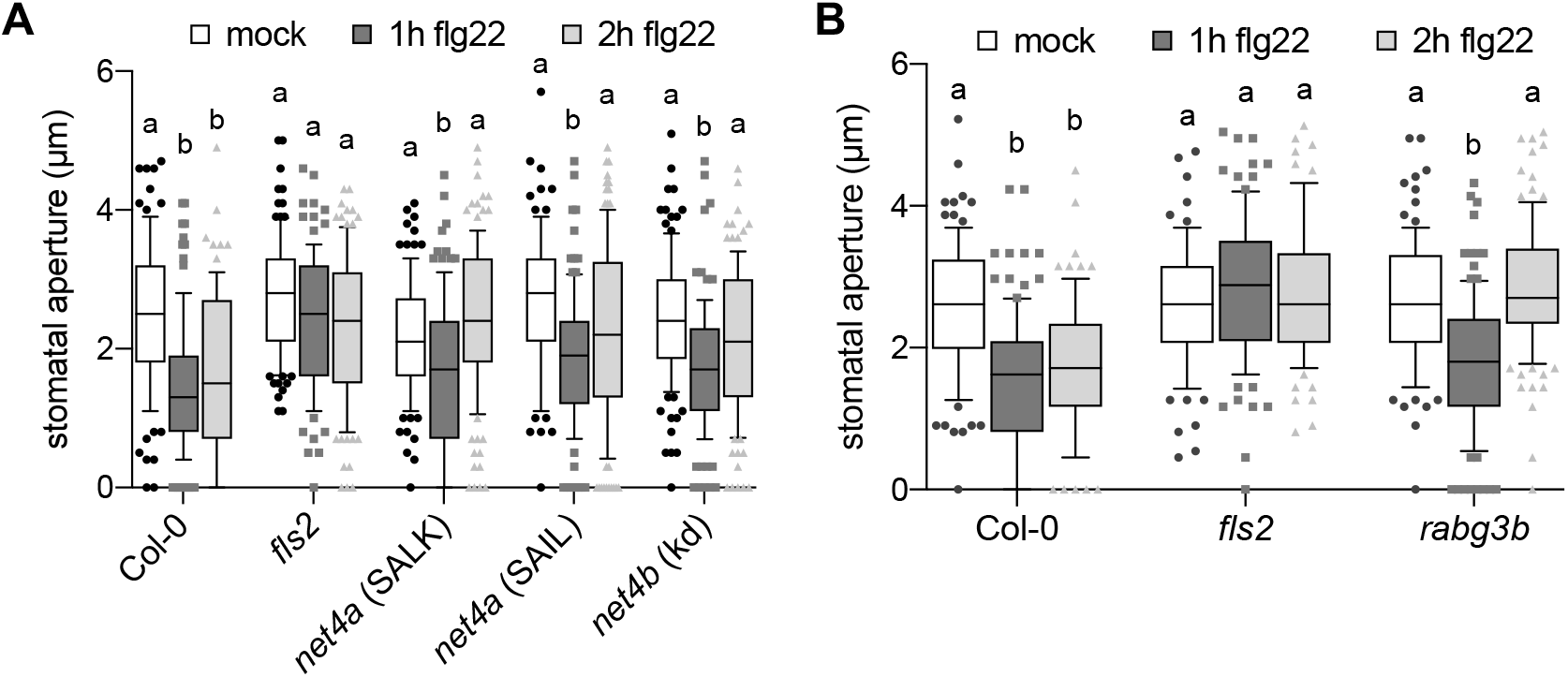
*net4* and *rabg3b* mutants are impaired in long stomatal closure but not generally immune-compromised. **(A, B)** Stomatal apertures measured in the indicated genotypes 1 and 2 hr after treatment with 20 μM flg22. Box plots of the values are shown with whiskers from the 5th to 95th percentiles, the line in the box shows the median (**(A)** n = 99-122 stomata (mock), n = 99-125 stomata (1 hr flg22), n = 91-125 stomata (2 hrs flg22); **(B)** n = 78-106 stomata (mock), n = 92-112 stomata (1 hr flg22), n = 86-116 stomata (2 hrs flg22)). Different letters indicate significantly different values at p < 0.0001 (2-way ANOVA, multiple comparisons). The experiment was repeated twice with similar results.

### NET4 proteins and RABG3 appear not to be involved in guard cell vacuole trafficking

Previous work demonstrated that RABG3 family members, together with RAB5 GTPases, mediate cargo trafficking to protein storage vacuoles ^26^. Signalling-active, flg22-bound FLS2 is degraded involving endocytosis and trafficking along RAB5 GTPase-positive endosomal compartments destined for lytic vacuoles ^35, 36, 37^. However, in FLS2 immunoblots we observed wild type-like flg22-reduced FLS2 abundance in *net4a* mutants (Fig. 3G), suggesting its normal degradation and endocytosis ^38, 35^.

RABG3b has been previously implemented in the regulation of autophagy related to pathogen-triggered and developmental cell death ^38, 39^. Also, autophagy has been linked to steady-state abundance of FLS2 and with stomatal movements ^40, 41^. Thus, we decided to test autophagic flux in *net4ab* double mutant and *rabg3b* upon flg22 treatment. Autophagic flux is defined as the rate of delivery of autophagic substrates to the vacuole and used as a measure to assess autophagy in different mutant backgrounds ^42^. We measured protein levels of (i) ATG8, the core autophagy protein that marks autophagosomes ^43^, (ii) NBR1, a well-established autophagy substrate ^44^, and (iii) catalase, a peroxisomal protein, since a recent study has shown that stomatal autophagy is crucial for peroxisome recycling ^41^. To assess vacuolar delivery, we treated cells with Concanamycin A, which inhibits vacuolar ATPase and leads to the stabilization of autophagic substrates ^44^. As a control, we used the *atg5* mutant, since in this mutant autophagic degradation is blocked ^45^. Our immunoblot analysis in wild type seedlings revealed a strong flg22-dependent autophagy induction 2 hrs after treatment and recovery (Fig. S4A-C). Autophagic flux experiments using these conditions showed no significant difference between wild type, *net4ab* and *rabg3b* mutants (Fig. S4D). Thus, despite we cannot exclude guard cell specific roles, as we analysed whole leaves and seedlings, we did not find major effects related to FLS2 trafficking and autophagic flux in *net4* and *rabg3b* mutants.

### Vacuole morphology appears unaltered in *net4* and *rabg3b*

A subcellular output of stomatal closure is the change in vacuole morphology ^12^, and recently it was shown that NET4 proteins modulate the compactness of vacuoles ^11^. Moreover, inhibition of RABG3 function in higher order mutants and by expression of a DN-RABG3f variant resulted in enlarged late endosomes and fragmented vacuoles ^26, 46^. To observe vacuole morphology in guard cells, we developed transgenic lines expressing the tonoplast marker YFP-VAMP711 in the *rabg3b* background ^47^. Confocal microscopy revealed no obvious effects in guard cell and pavement cell tonoplast morphology in *rabg3b* compared to wild type (Fig. S5A). Similar observations were made with *rabg3b* mutants expressing GFP-RABG3f (Fig. S5B). Flg22-stimulated wild type guard cells exhibited tonoplast patterns reminiscent of deflated vacuoles (Fig. S5C). Guard cells of *rabg3b* showed tonoplast patterns of inflated vacuoles independent of flg22 treatment (Fig. S5C).

We next used 2′, 7′-bis (2-Carboxyethyl)-5(6)-carboxyfluorescein (BCECF) as a luminar vacuole stain ^48^, revealing no difference in uptake between wild type and *rabg3b* guard cells and independent of the treatment conditions (Fig. S5D). Using 3D image reconstruction and surface rendering we measured guard cell vacuole volumes (Schmid et al., 2010) ^50^. Consistent with triggering stomatal closure, flg22-treated wild type guard cells showed a strong decrease in the vacuole volume and structures reminiscent of vacuole fragmentation or intra-vacuolar structure (Fig. S5E, S5F) ^12^. The volume of vacuoles in *net4ab* and *ragb3b* guard cells were not changed upon 2 hrs flg22 exposure, because in this condition stomata were open in these mutant backgrounds (Fig. S5D, S5F). These data show a correlation between vacuole morphology and the open and closed status of stomata ^12^, which is also the case for open stomata in 2 hrs flg22-treated *net4* and *rabg3b* mutants.

### NET4A/B affects the dynamic reorganization of the actin cytoskeleton

It was previously shown that PTI signalling changes the guard cell actin cytoskeleton organization ^23, 28^. Since NET4 proteins interact with F-actin and co-localize with actin filaments (Fig. 1, S1), we next investigated the effect of NET4A/B on cytoskeletal organization. We developed transgenic Col-0 and *net4ab* double mutant plants expressing the actin marker Lifeact-mNeon and applied quantitative image analysis of actin filament organization in guard cells from these lines. We measured the mean angular difference and parallelness of actin filaments ^28^, which were similar in guard cells of open stomata from Col-0 and net4ab (Fig. 7A-C). Interestingly, guard cells from *net4ab* mutants showed a lower value in the mean angular difference and a higher value in parallelness compared with Col-0 guard cells when treated with flg22 for 1 hr (Fig. 7D-F). At 2 hrs flg22 treatment, the mean angular difference and parallelness of actin filaments was similar in Col-0 and *net4ab* guard cells (Fig. 7G-I). Over the time points of 1 and 2hrs flg22 treatment, incremental changes occurred in Col-0 guard cells, showing the lowest value in mean angular difference and the highest value in parallelness at 2 hrs (Fig. 7A, 7D, 7G). This suggests that the actin cytoskeleton changes from a radial to a more longitudinal organization, a pattern correlating with stomatal closure and consistent with closed apertures in flg22-treated Col-0 (Fig. S6). By contrast, the mean angular difference was lowest and the parallelness was highest at the 1 hr time point in *net4ab* guard cells, which increased and decreased, respectively, at 2 hrs flg22 treatment (Fig. 7B, 7E, 7H). The latter suggests a recovery from a more longitudinal to a radial actin filament organization, which is correlated with guard cell opening. These results reveal that the *net4ab* mutants are compromised in guard cell actin cytoskeletal remodelling with an enhanced actin rearrangement, showing a stronger response at 1 hr and an earlier shift in the recovery at 2 hrs flg22 treatment. This demonstrates that actin cytoskeletal organization by NET4 proteins is required for normal stomatal closure to flg22, possibly inhibiting the reopening of stomatal pores.

**Figure 7.**
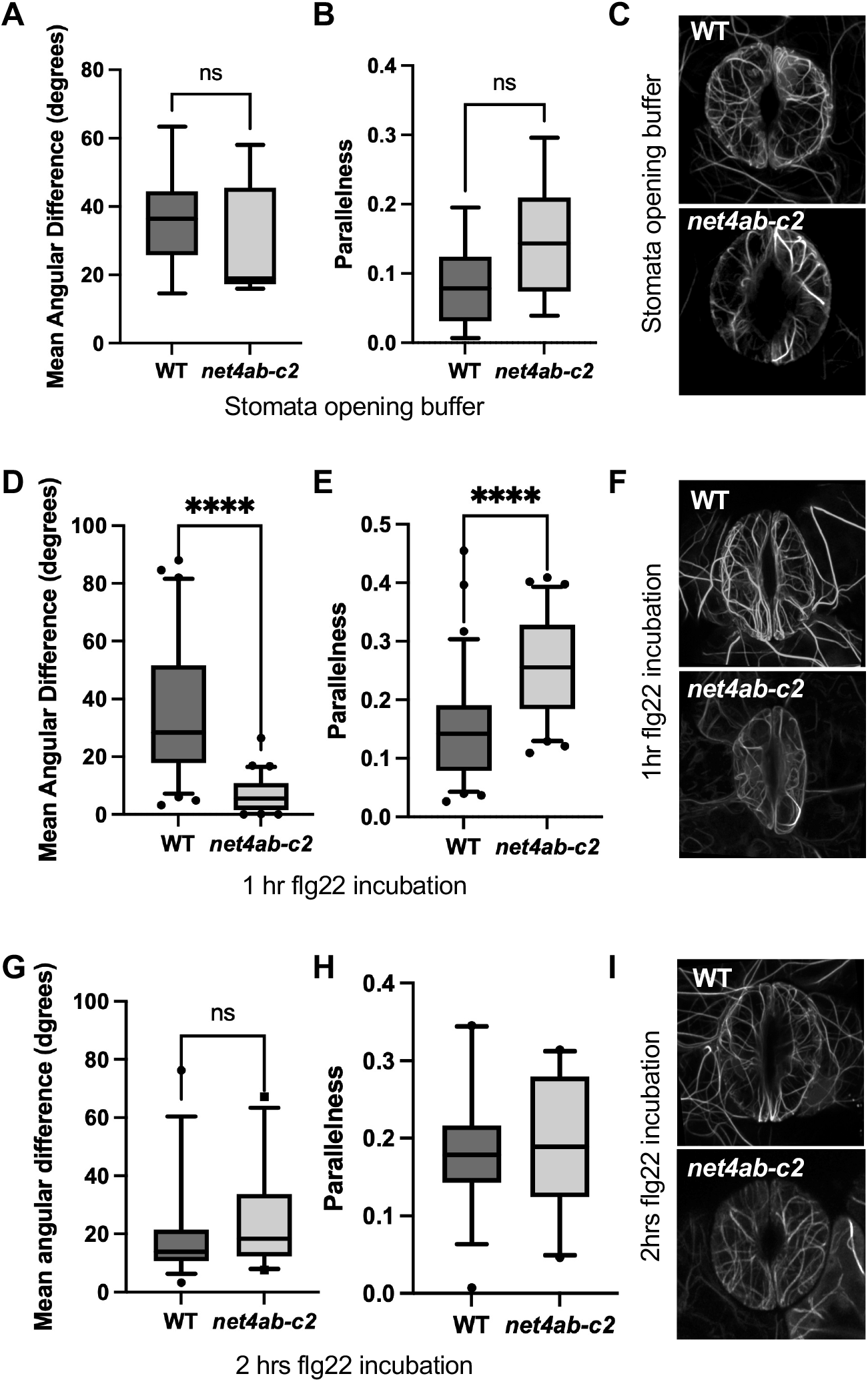
Guard cell actin dynamics are altered in *net4* mutants. **(A, B, D, E, G, H)** Quantification of actin cytoskeletal patterns of Lifeact-mNeonGreen-expressing Col-0 WT and *net4ab-c2* plants in guard cells. ImageJ analysis using the LPIXEL Inc LPX plugin set. **(A, B)** Mean angular difference **(A)** and parallelness **(B)** of the actin cytoskeleton in untreated guard cells. **(D, E)** Mean angular difference **(D)** and parallelness **(F)** of the actin cytoskeleton in guard cells treated for 1 hr with flg22. **(G, H)** Mean angular difference **(G)** and parallelness **(H)** of the actin cytoskeleton in guard cells treated for 2 hr with flg22. **(C, F, I)** Representative superresolution airyscan confocal images of Lifeact-mNeonGreen-positive actin filaments in Col-0 WT and *net4ab-c2* untreated and treated for 1 and 2 hrs with flg22, respectively; 1hr flg22 treatment n = 64-73 guard cells p < 0.0001 Mann Whitney t-test; 2hr flg22 treatment n = 23-38 guard cells p = 0.0777 ns Mann Whitney t-test; Stomata opening buffer 7-10 guard cells - angle p = 0.3638 parallelness p = 0.1088 ns Mann Whitney t-test.

## Discussion

Using molecular and cell biology approaches, we provide evidence for a molecular link between actin cytoskeletal organization and the vacuole morphologies underlying guard cell shape changes. We show that the two NET4 proteins have the potential to localize simultaneously to actin filaments and the vacuole tonoplast, mediated by two distinct domains: the family-characteristic NAB domain is involved in F-actin binding whereas the newly identified specific IRQ domain confers the direct interaction with RABG3 GTPases (Fig. 1, 2, 4, S1). This is compatible with the possibility that the tonoplast localization of NET4A and NET4B occurs through RABG3 GTPases. Indeed, proper NET4 localization depends on the presence of the IRQ domain and RABG3b co-expression (Fig. 5D). We noted that the interaction between the NET4 proteins and the RABG3 GTPases perfectly correlated with the tonoplast-localized and active, GTP-bound state of the Rab7 GTPases (Fig. 4) ^30^. A simple possibility to explain this correlation is to postulate that NET4A and NET4B are effectors of RABG3 GTPases. Consistent with this hypothesis is that *net4* and *rabg3b* mutants have similar phenotypes, exhibiting impaired stomatal closure after 2 hrs PAMP treatment (Fig. 5, 6). Thus, the tonoplast localization of the NET4 proteins can be linked to RABG3 GTPases, which may aid actin-tonoplast tethering in an activation-dependent manner.

Remodelling of the actin cytoskeletal organization is increasingly acknowledged as a critical function of plant immunity including stomatal movements ^51^. Moreover, patterns of actin cytoskeletal organization have been associated with different vacuole morphologies in plant growth ^10, 11^. In this study, we identify that the NET4 proteins are involved in actin remodelling that is triggered by FLS2 signalling. Mutant *net4* guard cells are compromised in actin filament remodelling, resulting in impaired long stomatal closure. Our results are compatible with a model, in which NET4 proteins participate in shifting actin cytoskeletal patterns (Fig. 7), a process coinciding with stomatal closure. Our data suggest that an excessive shift to a longitudinal actin filament pattern can result in closed stomata at 1 hr elicitation but the recovery to a more radial pattern already at 2 hrs compromises stomatal closure.

We did not observe global changes in guard cell vacuole volumes and morphologies (Fig S6). However, it is possible that the *net4* mutant guard cell vacuoles have defects in the formation of intra-vacuolar structures and thus have more compact vacuoles ^11, 12^. The function of the NET4-RABG3b interaction may be that of promoting actin-tonoplast contact site formation, thus mediating actin cytoskeletal remodelling along with vacuole morphology changes. NET4-RABG3b contact sites may help to properly align the actin filaments with the vacuole surface. During stomatal closure, actin filaments tethered to the tonoplast could reorganize into longitudinal arrays. This cytoskeletal remodelling could assist the deflation of the guard cell vacuole resulting in pore closure, a process driven by ion and water fluxes ^18^. Aberrant cytoskeletal remodelling would still allow initial guard cell vacuole deflation but impair the duration of the pore closure, as the recovery from an excessive longitudinal pattern would be earlier than normal. Thus, guard cell actin remodelling functions not only in the initial signalling-induced stomatal response but also in a longer lasting inhibition of reopening stomatal pores. Notably, our model would accommodate for the need of multiple cycles of remodelling/aligning, in that the activity of RABG3b may release the NET4 tethers from the interaction, allowing for reloading of the GTPase with GTP and further cycles. The signal-specificity and responsiveness of the *net4* mutant stomatal phenotype also fulfils the requirement for reversibility of the molecular and thus subcellular interactions in our model.

Activation of PRRs leads to the closure of stomata, through short signalling cascades regulating ion channels and ROS production from RBOHD ^18, 22, 52^. Active PRRs also induce actin cytoskeletal changes, involving the calcium-dependent protein kinase CPK3, downstream of FLS2, and its substrate ADF4 ^53, 54^. Future research will answer whether RABG3b could be a substrate of FLS2 signalling and upon phosphorylation interact with a GTP exchange factor, similar to human RAB7 tyrosine-phosphorylation by the Src kinase ^55^. If so, this signalling-dependence of additionally available GTP-bound, active RABG3b would further promote the process of NET4-driven vacuolar tethering of actin filaments potentially helping a longer lasting inhibition of reopening stomatal pores. Molecular links between actin-membrane interactions and signalling events have been identified previously between NET2A PRK4 and 5 ^5^, suggesting that short signalling cascades could regulate actin-membrane contact sites and therefore the underlying actin organisation.

The physiological significance of PAMP-induced stomatal closure and its molecular mechanisms are increasingly understood ^18^. Infectious pathogens actively interfere with the signalling events leading to pore closure or activate opening of stomata ^18^. Pathogens are also known to subvert host subcellular pathways, for example by targeting the actin cytoskeleton and Rab GTPases ^56, 57, 58, 59^, altogether highlighting a crucial role of actin- and Rab-dependent processes in immunity. Whether NET4 proteins and RABG3b are effector targets remains to be shown.

We note that *net4* and *rabg3b* mutants have no obvious global phenotypes. Also, the impaired stomatal closure is specific to PAMPs and no general phenotype. This raises questions whether other members of the NET family could have partial redundant functions to NET4A and NET4B, for example NET3A/C, also having IRQ domains (Fig. 4C). Roles for NET4 proteins beyond the guard cell may require distinct RabG isoforms found in those tissues and here, the additional IRQ domain containing NET3A/C may partially perform NET4 functions. Of note, guard cell immune responses appear to involve components not essential for other immune responses including the calcium-permeable channel OSCA1.3, which is also not required for ABA-induced stomatal closure ^60^.

Taken together, in this study we provide a molecular-subcellular framework of PAMP-induced stomatal closure through NET4-RABG3b. Our findings that NET4 proteins are involved in shifting actin filament reorganisation suggests that the inhibition of reopening stomatal pores is directed by actin cytoskeletal remodelling and its timing. Our findings also raise the possibility that RABG3b, which has already been implicated in membrane fusion at the vacuole ^26^, may have additional functions in actin rearrangements.

## Materials and Methods

### Plant materials and growth conditions

For assays with adult Arabidopsis plants, plants were grown on soil for 5 weeks in controlled environment rooms under short day conditions (10 h light at 22 °C and 14 h dark at 20 °C, 65% humidity). Arabidopsis seedlings were grown under sterile conditions in 24-well plates containing liquid medium (Murashige and Skoog Basal Medium (including vitamins), Duchefa, Haarlem, The Netherlands) for 14 d at 22 °C in a 16 h photoperiod. For live cell microscopy, transformed and crossed Arabidopsis lines were grown in the same conditions as described above. For *Nicotiana benthamiana* leaf transient transformation assays, *N. benthamiana* plants were grown on soil in controlled environment rooms under a 16 h photoperiod at 25 °C and 8 h darkness at 21 °C.

*fls2* mutants have been described previously 66. Homozygous T-DNA insertion lines *rabg3a-f, rabg2, net4a* and *net4b* were obtained from SAIL and SALK populations (Table S4). To constitutively express VAMP711-YFP in *rabg3b* mutant background, the pNIGEL07 vector carrying VAMP711-YFP fusion protein under control of the UBI10 promoter was transformed into SALK_004938 ^62^. Stable transgenic GFP-RABG3b lines were generated by cloning the genomic coding sequence of *RABG3b* either into the pUBN-GFP-Dest vector for constitutive expression or in fusion with 1180 bp of its native promoter into pCAMBIA1390 and Agrobacterium-mediated stable transformation into the *rabg3b* mutant background ^62, 63^. The FLS2-GFP construct was described previously ^61^. All constructs were confirmed by sequencing. *atg5-1* (AT5G17290, ATG5, SAIL_129B07) mutants have been described previously ^45^.

Stable transgenic *pNET4A*::NET4A-GFP lines were generated by cloning 2 kb of the promoter and genomic sequence of *NET4A* into pMDC107 and Agrobacterium-mediated stable transformation into Arabidopsis Col-0. *pNET4A*::NET4A-GFP lines were crossed with Ubi10::mCherry-RABG3f 50 for generating stable transgenic co-expressing lines. To create *NET4A* or *NET4B* promoter GUS reporter lines, 2kb promoter sequence of each gene was cloned in pBI101G and introduced into Col-0 by Agrobacterium-mediated stable transformation. Double *net4a/net4b* mutants were created by CRISPR/Cas9 gene editing. The *net4a*.*1* T-DNA insertion line was transformed with a vector driving EC1.2en:EC1.1 controlled expression of 3×FLAG-NLS-zCas9-NLS, which carried two *NET4B* gRNA modules. The nature of the mutation in the *net4ab* CRISPR-Cas9 double mutant is a t insertion in *NET4B*, which gives a frameshift in the latter part of the NAB domain, leading within two codons to a STOP codon. To analyse CRISPR/Cas9 generated genomic editing events, fragments surrounding the target sites were amplified by PCR using gene-specific primers N4B_F and N4B_R and sequenced using a nested primer (N4B_Fn1).

Actin marker lines expressing Lifeact-mNeonGreen were generated through Agrobacterium-mediated stable transformation of Col-0 and *net4ab* CRISPR-Cas9 double mutants. The Lifeact-mNeonGreen DNA coding sequence was synthesised (Integrated DNA Technologies) including gateway arms for downstream cloning. The final destination clone used to transform *net4ab* CRISPR-Cas9 double mutant consisted of this sequence recombined into pH7WG2.

### Transient transformation of *N. benthamiana*

For transient expression of proteins in *N. benthamiana*, constructs were transformed into *Agrobacterium tumefaciens* strain GV3101 and syringe infiltrated into young leaves. Cultures were infiltrated at OD_600_ between 0.1 and 0.4 and leaves imaged 2-3 days post infiltration. To transiently express NET4A and NET4B, their coding sequences were recombined into pCaMV35S driven binary GFP fusion destination vectors: pMDC43 (N-terminal GFP), pMDC83 (C-terminal GFP), pH7WGR2 (N-terminal mRFP). GFP-NET4BΔIRQ+CT was generated by cloning NET4B coding sequence residues 1-464 into pMDC43. mCherry-FABD2 construct encodes the C-terminal half of AtFIMBRIN1, which encompasses the second actin-binding domain of the protein (Voigt et al. 2005) fused to an N-terminal mCherry fluorescent tag. Lifeact-RFP consists of the 17 aa actin binding domain of Lifeact cloned into pH7RGW2 to produce an actin marker with a C-terminal red fluorescent protein tag ^5^.

### Stomatal closure assay

Leaf discs were harvested from at least 4 individual adult Arabidopsis plants per genotype and treatment using a biopsy puncher (4 mm width) and incubated for 3h in the light in stomata assay buffer (10mM 2-(N-morpholino)-ethanesulfonic acid (MES) pH 6.1, 50 mm KCl, 10 μm CaCl_2_, 0.01% Tween). After a subsequent 2 h treatment with 10 μM ABA or 20 μM flg22 (in stomata assay buffer in the light), stomata in the lower leaf epidermis were imaged using a bright field microscope DM-R (Leica, http://www.leica.com) at 40x magnification. Three images per leaf discs were captured and aperture sizes of the stomata within the field of imaging were measured using ImageJ ^64^.

### Bioassays for PAMP-induced responses

ROS assays were performed as described previously ^65^. Briefly, leaf discs were harvested with a biopsy puncher (4 mm width), placed in a 96-well plate, with a minimum of 12 leaf discs per genotype per assay, and incubated overnight in 200 μl of water in the dark. The water was then replaced with 100 μl of a luminol (0.4 mM)/ peroxidase (20 μg/ml^−1^ horseradish peroxidase, Sigma, www.sigmaaldrich.com) solution with or without (mock) 100 nM flg22. A Photek camera system and software were used to detect the luminescence over 45 min. Quantification of luminescence are binned values from 60 seconds.

### Gene expression analysis

Gene expression was studied in cDNA from whole seedlings, 15-days old vertically grown seedlings of the homozygous T-DNA insertion lines (*net4b*.*1, net4a*.*1* and *net4b*.*1/net4a*.*1*), and in cDNA from enriched guard cell and mesophyll extracts (obtained from H. Kollist; University of Tartu, Estonia) ^66^. Transcripts of the respective genes were quantified using real-time quantitative PCR with SYBR Green reagents and procedures (Applied Biosystems or SensiFAST SYBR DNA polymerase No-ROX Kit), and amplification was detected with the ABI PRISM 7700 Sequence Detection System or Rotor-gene Q PCR machine (Qiagen). For normalisation, the ΔCT method was used ^67^. Reverse transcription-PCR was also used to analyse the disruption of the *NET4B* and *NET4A* transcripts in the homozygous *net4b*.*1, net4a*.*1* and *net4b*.*1/net4a*.*1* T-DNA insertion lines. RT-PCR and qPCR primers are summarised in Table S5.

### GUS expression analysis

Plant tissue was incubated in GUS staining solution (100 mM phosphate buffer, 10 mM EDTA pH8.0, 0.1% (v/v) Triton X-100, 0.5 mM potassium ferricyanide, 0.5 mM potassium ferrocyanide, 1 mM X-Gluc (5-bromo-4-chloro-3-indolyl-β-D-glucuronide, Melford). Samples were imaged using a Zeiss Axioskop, Leica DM2500, and a Leica MC165 FC microscope with a Photometrics Coolsnap cf camera (Openlab 3.1.1 software) or Leica DFC420C camera (LAS AF).

### Yeast Two hybrid one-on-one interaction assay

Full length NET4A, NET4B and RAB coding sequences, and truncation fragments were Gateway-cloned into pGBKT7 or pGAD bait and pray vectors. CA- and DN-RABG3 variants were generated by site directed mutagenesis with QuickChange Lightning II Site-Directed Mutagenesis Kit (Agilent Technologies) according to manufacturers’ instructions. Bait and prey vectors (pGBKT7 and pGADT7) were transformed into the yeast strains AH109 and Y187, respectively. Single colonies of the bait and prey vector were mated on solid YPDA plates at room temperature for 24-48 hrs. As a negative control for auto-activation, above colonies were also mated with the respective empty vectors. Mated cells were selected by transfer onto to solid SD -WL dropout media. Diploid colonies were transferred onto selective SD dropout media -WLH + 2.5mM 3AT, and incubated at 30 °C for 5-7 days to test for potential interactions. As a positive control for yeast growth, diploid colonies were also grown on -WL SD plates. Yeast-two-hybrid screening using CA-RABG3b as a bait against a Arabidopsis prey cDNA library done by HYBRiGENiCS (https://www.hybrigenics-services.com/).

### Protein biochemistry

For co-immunoprecipitation (co-IP) from Arabidopsis, seedlings were grown in 6-well plates containing liquid MS medium (1% sucrose) and for each time point and genotype, seedlings of one 6-well plate with 15-20 seedlings per well were pooled and transferred into a beaker containing 50 ml liquid MS medium. After 1h of resting, flg22 (10 μM) was added to all samples but mock treatment and vacuum infiltrated for 5 min. Seedlings were harvested at the indicated time points and flash frozen in liquid nitrogen and subjected to protein extraction. Mock treatment was harvested at 30 min after treatment.

For co-IP from *Nicotiana benthamiana* plants, 5-6 weeks old plants were infiltrated with Agrobacteria strains (GV3101 pMP90, OD_600_ 0.3) expressing the proteins of interest. After 3 days, 2 infiltrated leaves were detached and transferred into a beaker containing 150 ml H_2_O supplemented with 10 μM flg22 or not (mock), which was vacuum infiltrated for 3 min. Leaves were dried and flash frozen in liquid nitrogen. Tissue was grinded and transferred to centrifugation tubes. Total proteins were extracted using 1 ml of extraction buffer (50 mM Tris HCL pH 7.5, 50 mM NaCl, 10% glycerol, 2 mM EDTA, 2 mM DTT, 1% protease inhibitor cocktail (SIGMA), 1% PMSF, 1% phosphatase inhibitor cocktails 2 and 3 (SIGMA), 1% IGEPAL) per 1 ml grinded plant tissue. Samples were incubated in extraction buffer for 30 min rotating at 4 °C. Upon centrifugation (20 min, 15000 rpm, 4 °C), supernatant was filtered with miracloth and BioRad column. A 100 μl aliquot of the supernatant was supplemented with 30 μL 6xSDS sample buffer and used as input protein sample. 10 μl GFP-Trap®_A beads (chromotek; www.chromotek.com) were added to the residual supernatant and the immunoprecipitation incubated for 3 h rotating at 4 °C. Beads were collected upon centrifugation (1min, 500xg) and washed three times in wash buffer (50 mM Tris-HCl pH 7.1, 50 mM NaCl, 1% PMSF, 0.5% IGEPAL). After the final wash, the supernatant was completely removed and 60 μl 2x SDS sample buffer was added to the beads (IP). Before loading and running 25 μl of the samples on an SDS PAGE (10% SDS gel), they were boiled for 10 min at 95 °C. Proteins were transferred from the SDS gel onto a PVDF membrane through semi-dry western blotting. Membranes were incubated with the respective primary antibodies as indicated. After 3 washes with TBS-T, membranes were incubated with the respective secondary antibodies for 1.5 h. After washing the membranes in TBS-T, membranes were developed using SuperSignal™ West Femto Maximum Sensitivity Substrate (Thermo Fisher) and exposure to X-ray film (Fujifilm Super RX) for 2 min to 4 h, depending on the samples.

Tandem affinity purification coupled to mass spectrometry (TAP-MS) of TAP-tagged NET4A was carried out as part of the AP-MS platform service provided by the Functional Interactomics group at the Centre for Plant Systems Biology VIB-Ghent. The NET4A ORF was transferred from pDONR207 into a GSrhIno tag vector by Gateway cloning to fuse with a double affinity tag, which consists of a protein G tag and a streptavidin-binding peptide separated by a very specific protease cleavage site. Arabidopsis cell suspension cultures were transformed, and *in planta* protein complexes were isolated using two sequential purification steps and identified through LC-MS/MS.

To test for RABG3 interactors *in vivo*, total *Arabidopsis* root microsomes were isolated from plants expressing YFP::RabG3f (Wave-5Y) 50, YFP::RABA2a 74, YFP::RABA5c 75 or no transgene. 20-30 surface-sterilised seeds were grown in 20 ml medium (0.3x Murashige and Skoog medium (Sigma, Poole, UK), 1% sucrose, 0.05% MES pH5.7) under 60 rpm shaking, 16 h days and 21°C for 14 days. 3 days after addition of 5 μM naphthyl acetic acid, 4-5g roots were ground in extraction buffer (50mM HEPES pH7.5, 50mM NaCl, 5mM MgCl_2_, 10% glycerol, 0.5mM PMSF, 100μM GTP-□-S and Complete EDTA-Free Protease Inhibitor (Roche), centrifuged and 1 mM DTSSP (3,3’-dithiobis-(sulfosuccinimidyl propionate)) was added to the supernatant, incubated on ice for 30 min, followed by quenching with Tris-HCl pH7.5 at 50mM on ice for 10 minutes. Microsomes were pelleted onto a cushion of 2 M sucrose by centrifugation at 541,000x*g* for 30 min at 4°C and re-suspended in 1.8 ml extraction buffer with 2% CHAPS, and centrifuged at 541,000x *g* for 30 min at 4°C. 200 μl anti-GFP μMACS magnetic microbeads (Miltenyi Biotec) were added to the supernatant and incubated for 60 min on ice with gentle agitation. Bound proteins were eluted in 50 μl Laemmli protein loading buffer. Samples were separated on a NuPAGE Novex 4-12 Bis Tris polyacrylamide gel (Thermofisher Ltd). In-gel trypsin digestion and mass spectrometry were performed at the Central Proteomics Facility, University of Oxford (www.proteomics.ox.ac.uk) and proteins were quantified using the label-free SInQ pipeline ^69^. Co-immunoprecipitated proteins were quantified from at least 3 independent experiments and ranked according to their abundance in the YFP::RABG3f immunopreciptate relative to immunopreciptates with YFP::RABA2a and YFP::RABA5c and their abundance in total microsomes.

### MAPK assays

Two weeks-old seedlings (n=12) were placed in dH_2_O for 16 h. 1 μM flg22 was added for 10 min and tissue (50 mg per sample) was shock frozen. To the grinded material 50 μL SDS-PAGE sample loading (0.35M Tris-HCl pH 6.8; 30% [v/v] glycerol; 10% [v/v] SDS; 0.6M dithiothreitol; and 0.012% [w/v] bromphenol blue) was added. Total proteins were separated by electrophoresis in 12 % SDS-polyacrylamide gel and electrophoretically transferred to a polyvinylidene fluoride membrane according to the manufacturer’s instructions (Bio-Rad). Transferred proteins were detected with Ponceau-S. Polyclonal primary antibodies against phospho-p44/42 MAPK (Cell Signaling Technologies) were used, with alkaline phosphatase-conjugated anti-rabbit as secondary antibodies. Signal detection was performed using CDPstar (Roche).

### Autophagy assays

To investigate RABG3b, NET4A and NET4B involvement in autophagy, whole seedlings were grown in liquid ½ MS media (1% sucrose) in 12-well plates under continuous light and constant shaking at 80 rpm. For each genotype, treatment and time point 20 seedlings were grown. After 1 week, seedlings, except for mock, were treated with 10 *μ*M flg22 for either 1 h or 2 h. Afterwards the first round of samples was harvested. The other set of samples underwent a recovery period of 8 h in ½ MS media containing 1 *μ*M concanamycin A (conA) (CAS 80890-47-7; Santa Cruz), which has been dissolved in DMSO as a 2 mM stock solution, or DMSO (as control, same volume as conA). Since conA is light sensitive, seedlings (with and without conA) were subjected to dark treatment where plates were wrapped in aluminum foil. For all treatments, wells were first washed with 1 ml media containing respective treatment before incubation. Seedlings were harvested in microcentrifugation tubes containing a variety of different sized glass beads (2.85-3.45 mm, 1.7-2.1 mm and 0.75-1.00 mm; Lactan GmbH) and flash frozen in liquid nitrogen. Plant tissue was grinded with mixer mill MM400 (3x 30 s, 30 Hz; Retsch) and the proteins were extracted in 250 *μ*l extraction buffer containing 100 mM Tris, 200 mM NaCl, 1 mM EDTA, 2% 2-Mercaptoethanol, 0,2% Triton X-100 and cOmplete, EDTA-free Protease Inhibitor Cocktail (SIGMA) with a pH of 7.8. Well mixed samples were centrifuged at 15000 rpm for 10 min at 4°C. The whole supernatant was transferred and centrifuged again. 100 *μ*l of resulting supernatant was eluted in 2x Laemmli protein loading buffer and boiled for 10 min at 95°C. Protein amount was quantified via Amido black77. 10 *μ*l of sample was diluted in 190 *μ*l water and 1 ml of Amido black staining solution (90% methanol, 10% acetic acid, 0.005% (w/v) Amido black 10B (SIGMA)) was added. For the staining solution a Bovine Serum Albumin (BSA) standard curve was determined. Samples were mixed thoroughly, centrifuged for 10 min at 15000 rpm and the supernatant was discarded. Pellets were washed in 1 ml washing solution (90% ethanol and 10% acetic acid) and centrifuged again. After full removal of supernatant, samples were dissolved in 1 ml 0.2 N NaOH and 200 *μ*l was loaded on 96-well plates suitable for a plate reader (Synergy™ HTX Multi-Mode Microplate Reader; BioTek). Concentration was measured at 630 nm and calculated via the previously determined BSA standard curve where y stands for the measured optical density (OD) and x for the protein concentration. 15 *μ*g of protein was loaded on 4-20% Mini-PROTEAN® TGX™ precast gel (Bio-Rad) and blotted on nitrocellulose membrane using the semi-dry Trans-Blot® Turbo™ Transfer System (Bio-Rad). Total protein extract was immunoblotted with either anti-NBR1 (1:2000; Agrisera), anti-Catalase (1:10000; Agrisera) or anti-ATG8a-i (1:4000; Agrisera) antibodies. Images were acquired by developing with SuperSignal™ West Pico PLUS Chemiluminescent Substrate (Thermo Fischer) and detected via ChemiDoc™ Touch Imaging System (Bio-Rad).

### Actin co-sedimentation assays

The NET4B 1-105 aa NAB domain was Gateway-cloned into expression vector pGAT4, expressed as a 6xHis-tagged fusion protein in *E. coli* strain Rosetta 2 (DE3) pLysS (Novagen) and purified using Ni-NTA resin (Qiagen) according to manufacturer"s instructions. Recombinant proteins were dialysed into reaction buffer (4 mM Tris pH 8.0, 0.2 mM DTT, 0.4 mM ATP, 20 mM KCl, 4 mM imidazole, 2 mM EGTA, 0.4 mM MgSO4). Rabbit muscle actin (Cytoskeleton Inc.) in G-buffer (2 mM Tris pH 8.0, 0.5 mM DTT, 0.2 mM CaCl2, 0.2 mM ATP) was polymerized by the addition of 10 x KME (500 mM KCl, 10 mM MgSO4, 10 mM EGTA, 100 mM imidazole, pH 6.5). Actin (5um) and NET4B (10 μM) were mixed in reaction buffer at room temperature for 1 hr and then centrifuged at 350,000 x g for 15 minutes before supernatant and pellet fractions were compared using SDS-PAGE.

### Confocal microscopy

Confocal microscopy was performed on either a Zeiss 880 LSM with Airyscan detector module a, or Leica SP5 TCS confocal laser scanning microscope equipped with a 63x water or oil immersion objective. Leaf or root samples were mounted in water. Large root areas were acquired in tile scan mode with auto stitching. For *N. benthamiana* imaging, small sections were excised from the leaf and mounted on slides in water. Roots of stable Arabidopsis lines were mounted on slides in ½ MS with the green tissues free outside the coverslip chamber. Where multiple fluorescent proteins or markers were co-imaged, data was acquired sequentially in multitrack or multichannel mode with laser line switching. GFP and BCECF were excited using the 488-nm argon laser, and fluorescence emission captured between 500 and 550 nm for GFP, between 580 and 620 nm for FM4-64 and between 510 and 550 nm for BCECF. RFP was excited at 561 nm, and emission captured between 580 and 620 nm. mCherry was excited at 594 and emission captured 600-650 nm.

Forster resonance energy transfer-fluorescence lifetime imaging microscopy (FRET-FLIM) experiments were performed on a Leica SP5 TCS SMD confocal microscope with a Picoquant LSM FLIM upgrade kit. *N. benthamiana* leaves were transiently transformed by Agrobacterium infiltration with both NET4A-GFP and NET4B-RFP constructs. FLIM experimentation and analysis using PicoQuant SymPhoTime 32 software was carried as described in ^71^. All measurements were taken from whole-field images of cells expressing fluorophore fusion proteins at similar level, and at least 6 measurements were taken for each analysis.

### Actin imaging and quantification

Leaf discs (approx. 4mm width) of 4 weeks-old soil grown plants were submerged in stomata opening buffer (10mM 2-(N-morpholino)-ethanesulfonic acid (MES) pH 6.1, 50 mm KCl, 10 μm CaCl2, 0.01% Tween) and incubated in the light for 2h. Subsequently, the leaf discs were transferred into stomata assay buffer supplemented with 20 μm flg22, and incubated in the light for a further 1 or 2 hrs before imaging. Super-resolution Lifeact labelled actin arrays within guard cells were imaged using a Zeiss 880 LSM with Airyscan detector module (Excitation 488nm, emission 495nm-550nm). The actin orientation within z stacks (Mean angular difference and Parallelness) was quantified in ImageJ using the LPIXEL Inc (https://lpixel.net/en/products/lpixel-imagej-plugins/) set analysis of tools as previously described ^72 73 28 13^. Briefly, a maximum intensity projection was created from each Z stack which was then rotated so that the long axis of the guard cell was vertical. The maximum intensity projection of the stoma was then separated into individual guard cells. Using the freehand selection tool, the guard cell was outlined to create an ROI, which was subsequently added to the ROI manager. The cell medial axis (angle) was calculated in the ROI manager for each guard (using the “Fit Ellipse” option under “Set Measurements.”) The LPX Filter2d plugin, using the filter “lineFilters” and the linemode “lineExtract,” was used to skeletonize actin in each guard cell. The default settings for “lineExtract” were used (giwslter = 5, mdnmsLen = 15, shaveLen = 5, delLen = 5, preGauss = -1). Skeletonised filaments from surrounding cells were excluded from analysis by loading the guard cell ROI and using “clear outside”. The average theta and parallelness for each guard cell were calculated using the LPX Filter2d plugin, using the filter “lineFilters” and the linemode “lineFeature.” The mean angular difference was calculated by subtracting the angle of the cell medial axis over the horizontal (close to 90°) from the average theta and taking the absolute value.

### BCECF staining

To stain the vacuolar lumen of *A. thaliana* guard cells, the pH-sensitive dye BCECF-AM (Molecular Probes) was used. BCECF was prepared as a 1 mg/mL stock solution (∼1.67 mM) in DMSO. Leaf discs (4mm width) of 3 weeks-old soil grown plants were submerged in stomata assay buffer (10mM 2-(N-morpholino)-ethanesulfonic acid (MES) pH 6.1, 50 mm KCl, 10 μm CaCl2, 0.01% Tween) and incubated in the light for 2h. Subsequently, the leaf discs were transferred into stomata assay buffer supplemented with the respective elicitors, and incubated in the light for further 2 h. 90 min into the treatment, BCECF stock solution was added to a final concentration of 10 μM. After 30min of dye loading the leaf discs were imaged using a confocal microscope, leaf discs were washed twice in stomata assay buffer before imaging. Airyscan confocal z stacks of guard cell vacuoles loaded with BCECF were surface rendered and their volume quantified using Fiji with 3D Viewer plugin ^49, 50^. Volumes were also confirmed by segmentation and volume quantification with Amira for Life & Biomedical Sciences (Thermo Scientific).

### Immunogold labelling and transmission electron microscopy

A NET4B fragment (residues 121-220) was expressed as a His-tagged, recombinant protein and purified using Ni-NTA resin (Qiagen). Polyclonal antibodies were raised in mice as described in ^1^. Distal 1-2 mm root tips of 7 days-old Arabidopsis Col-0 seedlings were excised and immersed in 20% (w/v) BSA, and immediately high-pressure frozen with a Leica EMPACT high-pressure freezer. Samples were fixed by freeze substitution with a Leica EM automated freeze substitution (AFS) device (Leica Microsystems GmbH) using anhydrous acetone containing 0.25% (v/v) glutaraldehyde and 0.1% (w/v) uranyl acetate. Resin was gradually infiltrated (Monostep Lowicryl HM20 Agar Scientific) replacing acetone, polymerised and ultrathin sections (50-70 nm) prepared. Sections were rinsed with 0.1% (v/v) glycine PBS, blocked for 30 minutes in 1% (w/v) BSA PBS and finally incubated in 0.1% BSA-c (Aurion) PBS. Sections were incubated for 30 min with the primary antibody (rabbit anti-NET4B) or the pre-bleed sera as a negative control. Grids were washed in 0.1% BSA-c and then incubated for 30 minutes with the 5 nm colloidal gold-conjugated goat anti-rabbit secondary antibody (British Biocell International, Cardiff, UK). Sections were also incubated without the secondary antibody as a negative control for immunogold labelling (secondary omission control). The grids were rinsed in PBS and the antigen-antibody-gold complex stabilised by incubation in 1% (v/v) glutaraldehyde in PBS. Sections were imaged using a Hitachi H-7600 TEM operating at 100 kV fitted with an AMT Orca-ER digital camera (Advanced Microscopy Techniques, Danvers, USA). Quantification of tonoplast labelling was carried out using the Relative Labelling Index (RLI) method (Mayhew, Lucocq and Griffiths, 2002) (Mayhew and Lucocq, 2008). The tonoplast demonstrated the most abundant anti-NET4B labelling satisfying both criteria for preferential labelling giving a RLI of 3.58, and a Chi squared 70.34% of total.

## Author contributions

PJH and SR conceived the project and supervised the research. T.J.H., M.Ko., D.A.M, P.D., J.T.M.K., M.T., A.S., G.DJ., M.Ka., I.M., Y.D., P.J.H., and S.R. designed the methodology; T.J.H., M.Ko., D.A.M., P.D., J.T.M.K., M.Th., C.R., M.Te., A.S., G.DJ., M.Ka. performed research; T.J.H., M.Ko., D.A.M., P.D., J.T.M.K., M.Th., C.R., M.Te., A.S., G.DJ., M.Ka, I.M., Y.D., P.J.H., and S.R. analysed data; S.R. and T.J.H. wrote the paper with edits from M.Ko. and P.J.H.

## Acknowledgements

We like to thank members of the Robatzek and Hussey laboratories for fruitful discussions, Prof. Hannes Kollist (University of Tartu, Estonia) for providing materials, and Dr. Philip Charles (University of Oxford) for help with proteomics analysis.

