## Supplemental Information for "A NET4-RabG3 couple mediate the link between actin and the tonoplast and is essential for normal actin cytoskeletal remodelling in stomatal closure to flg22"

**Supplemental Information Hawkins and Kopischke et al.**

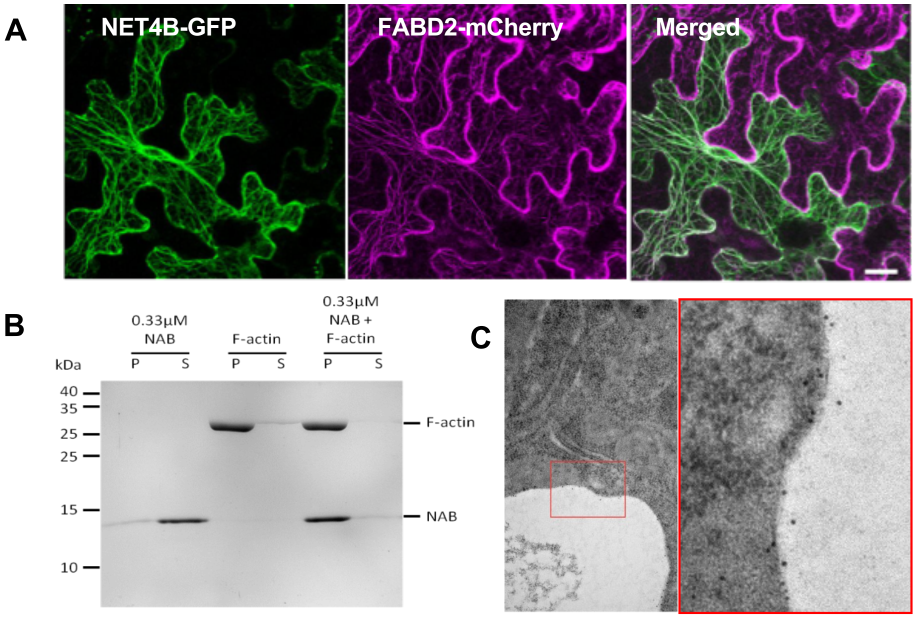


**Figure S1. NET4 proteins bind actin and localize to the tonoplast. (A)** Confocal microscopy of *N. benthamiana* leaves transiently co-expressing NET4A-GFP and the actin marker mCherry-FABD2. Representative images are shown; scale bar = 20 µm. **(B)** Actin co-sedimention assay. Recombinant NAB domains of NET4B are mixed and co-sedimented with F-actin following ultra-centrifugation. **(C)** Transmission electron micrograph of anti-NET4B immunogold-labelled root sections. Gold particles are located in the vicinity of the tonoplast membrane. The tonoplast demonstrated the most abundant anti-NET4B labelling satisfying both criteria for preferential labelling giving a RLI (Relative Labelling Index) of 3.58, and a Chi squared 70.34% of the total.

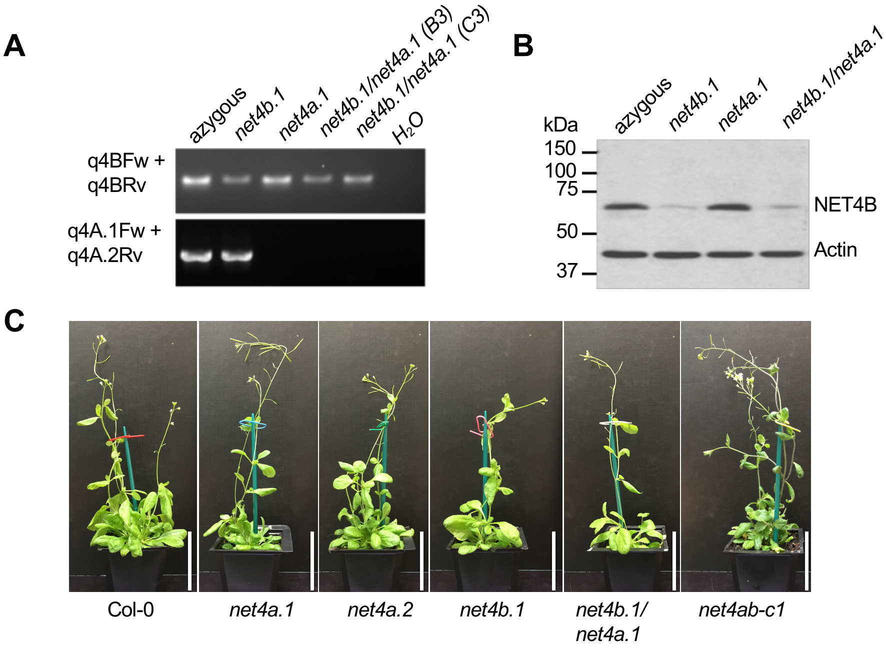


**Figure S2. Characterization of *net4* mutants and complementation lines. (A)** Transcript levels of *NET4A* and *NET4B* upon their genetic disruption in homozygous *net4b.1*, *net4a.1* and *net4b.1/net4a.1* T-DNA lines. No functional *NET4A* transcript is present in *net4a.1* and *net4b.1/net4a.1*. However, *NET4B* transcripts are present in *net4b.1* and *net4b.1/net4a.1* albeit at reduced levels, which represents a knock-down. **(B)** Immunoblot analysis of total protein extracts from *net4b.1, net4a.1, net4b.1/net4a.1*, and azygous plants using anti-NET4B antibodies. Blots show reduced NET4B protein levels in *net4b.1* and *net4b.1/net4a.1*. **(C)** Macroscopic analysis of *net4* mutants indicate normal plant development and growth.

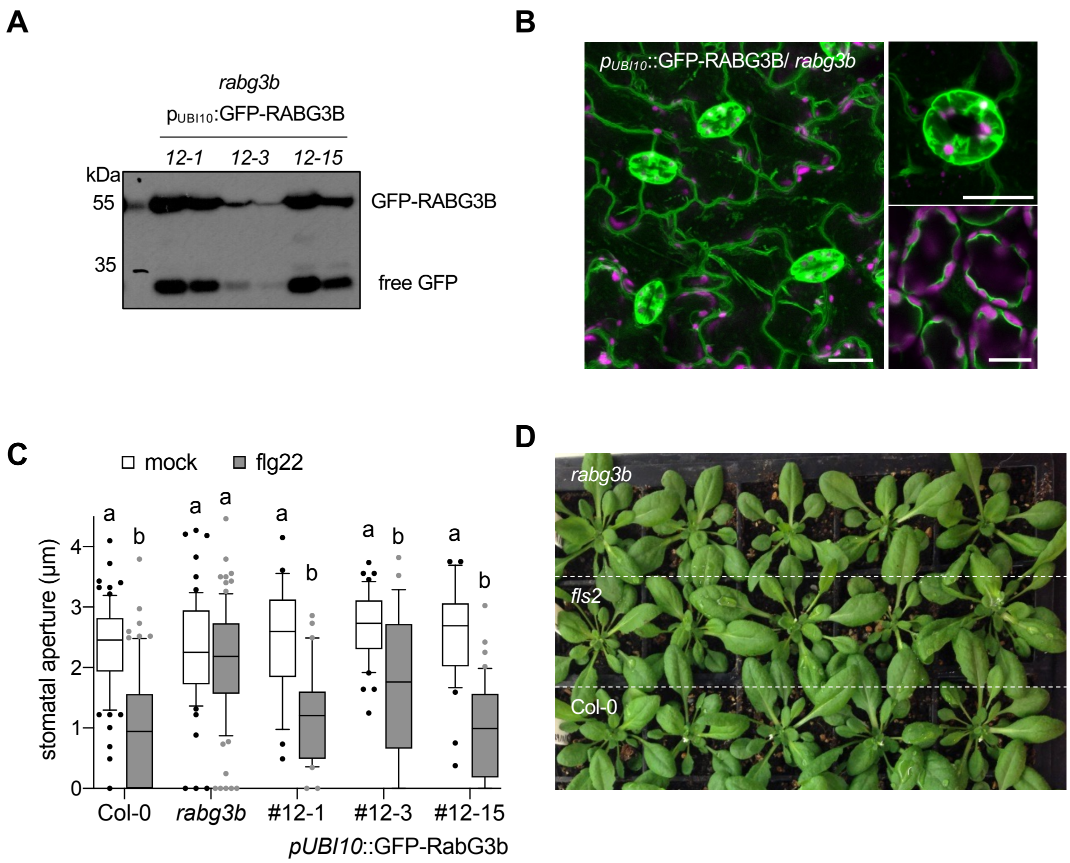


**Figure S3. Characterization of *rabg3b* mutants and complementation lines**. **(A)** Immunoblot analysis of *rabg3b/*GFP-RABG3b expressing lines. Full-length GFP-RABG3b was detected using anti-GFP antibodies. The level of mutant complementation was correlated with the expression level of tagged RABG3b. **(B)** Confocal micrographs of *rabg3b/*GFP-RABG3b expressing lines. GFP fluorescence at the tonoplast could be observed in epidermal, guard and mesophyll cells. Shown are representative images overlaying signals from GFP fluorescence (green) and chlorophyll auto-fluorescence (magenta); scale bars = 20 µm. **(C)** Stomatal aperture measurements in the indicated GFP-RABG3b expressing genotypes compared with Col-0 wild type and *rabg3b*. Stomatal apertures were measured 2 hr after treatment with 20 µM flg22. Box plots of the values are shown with whiskers from the 5th to 95th percentiles, the line in the box shows the median (n = 22-77 stomata (mock), n = 30-88 stomata (flg22)). Different letters indicate significantly different values at $p < 0.0001$ (2-way ANOVA, multiple comparisons). **(D)** Macroscopic analysis of *rabg3b* mutants indicate normal plant development and growth.

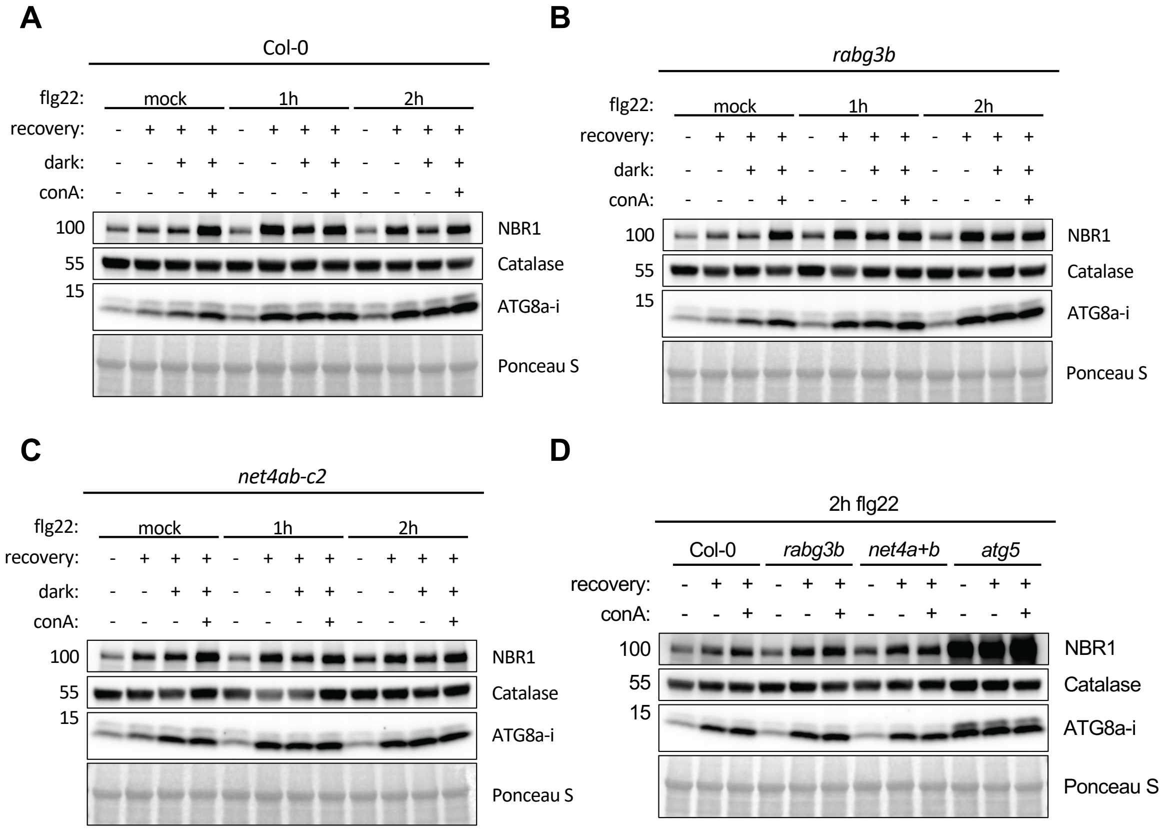


**Figure S4. Time course autophagic flux experiment for Col-0, *rabg3b* and *net4a/b* double mutant upon *flg22* treatment.** Whole seedlings were treated with 10 µM *flg22* for the indicated time points and underwent a recovery period of 8h in (i) light, (ii) dark or (iii) dark and 1 µM conA. 15 µg of protein was loaded on each lane. Proteins were detected via anti-NBR1, anti-Catalase and anti-ATG8a-i antibodies, respectively.

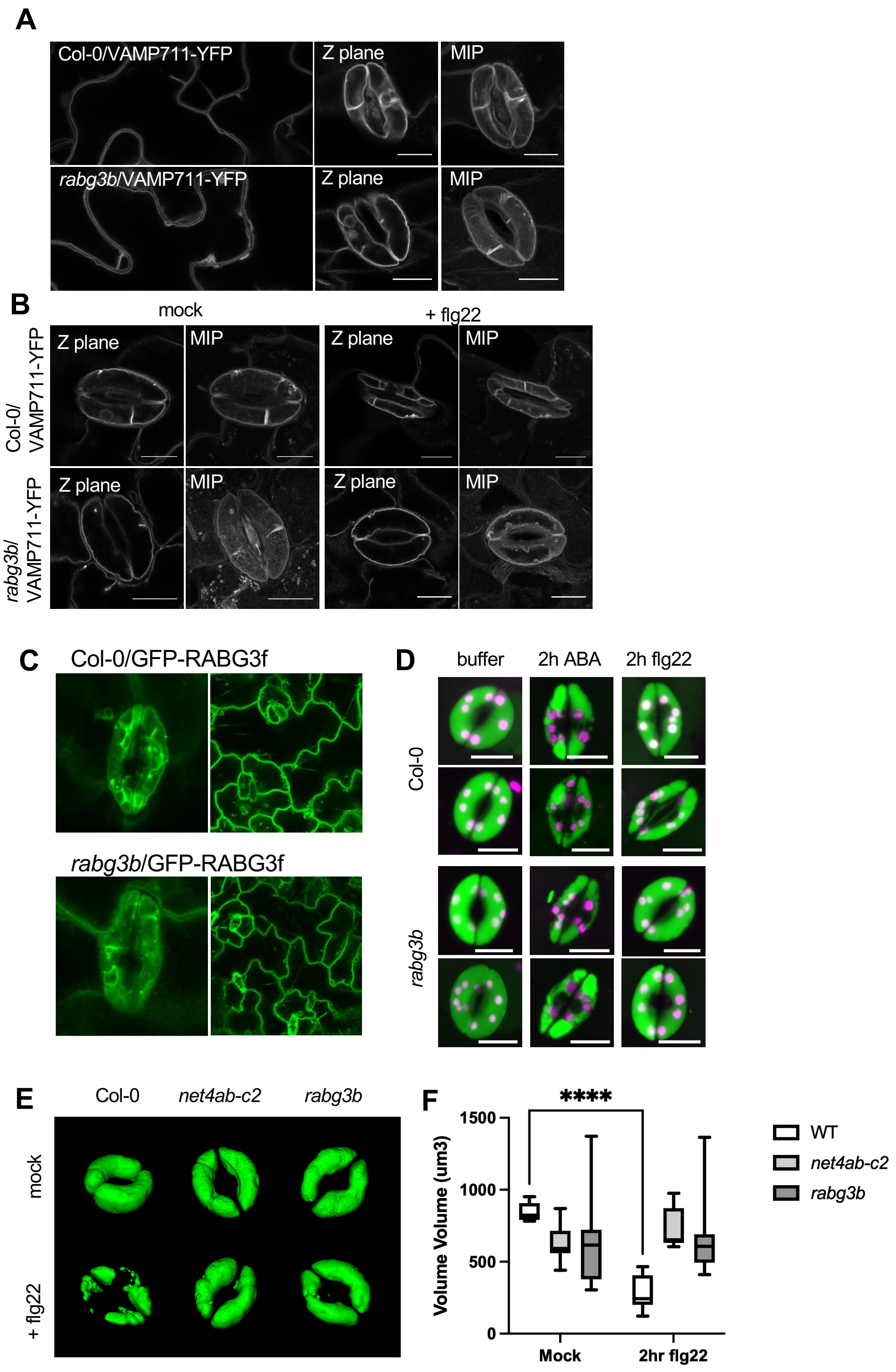


**Figure S5. Tonoplast morphology and vacuolar trafficking in *rabg3b* and *net4* mutants.**
**(A, B)** Confocal micrographs of tonoplast membranes in transgenic *rabg3b* mutant lines expressing the tonoplast marker YFP-VAMP711. **(A)** Tonoplast localization of YFP-VAMP711 in *rabg3b* pavement and guard cells were similar to wild type (Col-0). **(B)** Tonoplast localization of YFP-VAMP711 in *rabg3b* guard cells were similar to wild type (Col-0) whether open or upon closure induced by 20 µM flg22 for 2

hrs; scale bars = 10 µm (MIP = maximum intensity projection). **(C)** Confocal micrographs of vacuoles in leaf epidermis and guard cells of Col-0 (WT) and *rabg3b* seedlings visualised by constitutive expression of the tonoplast marker YFP-RABG3F. **(D)** Vacuolar lumen staining with BCECF. The BCECF staining was performed after treatment with ABA (10uM) or flg22 (20 µM) to observe vacuolar re-arrangements during stomatal closure. **(E, F)** Confocal microscopy of Arabidopsis guard cell vacuoles of the indicated genotypes treated with 20 µM flg22. Vacuolar lumen was stained with BCECF. **(E)** Representative images of 3D vacuole reconstructions are shown. **(F)** Measurements of volumes from 3D vacuole reconstructions. Bars represent mean values ± SEM of 3 biological replicates; n = 65 guard cells; ANOVA with Tukey's multiple comparisons test. Different letters indicate significantly different values at $p < 0.05$.

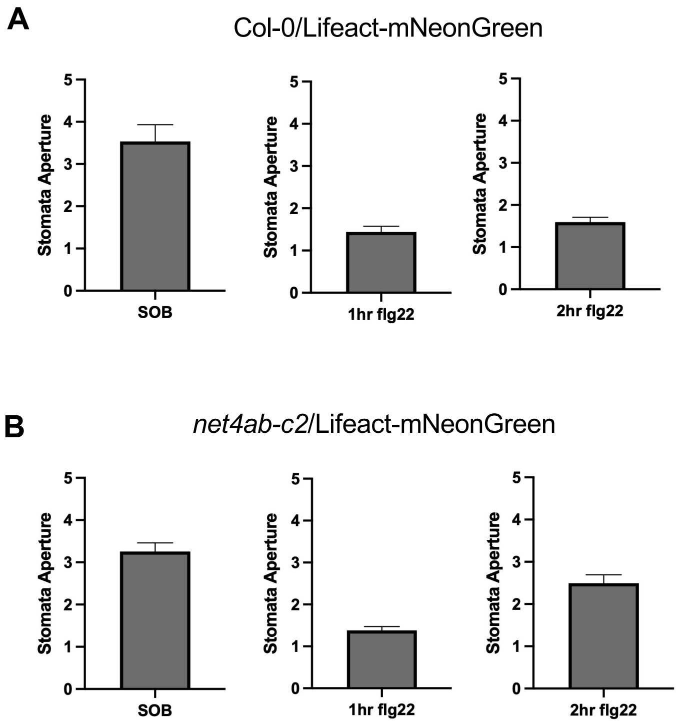


**Figure S6. Stomatal movements in Lifeact-mNeonGreen expressing Col-0 and *net4ab-c2* (A, B)** Stomatal aperture measurements of images used for quantitative evaluation of actin filament organization in guard cells from Col-0/Lifeact-mNeonGreen (WT) and *net4ab-c2*/Lifeact-mNeonGreen plants in stomata opening buffer (SOB) and after 1 and 2 hrs flg22 treatment.

**Table S1. Potential NET4A interactors retrieved from tandem affinity purification (TAP)-tagging.**

| **ATG Code** | | **1 % DDM** | **1% Digitonin** | **1% C12E8** | **1% TritonX100** | **Total** |
|---|---|---|---|---|---|---|
| AT5G58320 | AtNET4A | + | + | + | + | 4 |
| AT4G09720 | AtRABG3A | | + | + | + | 3 |

**Table S2. YFP-RABG3f interactors retrieved by immuno-precipitation from Arabidopsis.** Co-immunoprecipitated proteins were quantified from at least three independent experiments and ranked according to their abundance in the YFP::RABG3f immunopreciptate relative to immunopreciptates with YFP::RABA2a and YFP::RABA5c and their abundance in total microsomes. The most highly ranked protein was AtVPS26A, a known interactor of RABG3f (Zelazny et al., 2013). NET4B consistently co-precipitated with YFP::RABG3f but never with YFP::RABA2a or YFP::RABA5c, nor was it detectable in total microsomes (P(plgem) = 0.005, 0.018, 0.028 respectively).

| | | | Relative abundance* | | | | Peptide detection stats | | | | | |
|---|---|---|---|---|---|---|---|---|---|---|---|---|
| Protein | ID | Description | RAB-G3f | RAB-A2a | RAB-A5c | Total microsomes | Length (AA) | mw (Da) | Indistinguishable Proteins | PSMs | Peptide Seqs | % Coverage |
| VPS26A | sp\|Q9FJD0\|VP26A_ARATH | | 50.9 | 0.00 | 0.00 | 0.00 | 302 | 35182 | None | 69 | 30 | 87 |
| | | | 24.40 | 0.00 | 0.00 | 0.00 | | | | | | |
| P(plgem) | NH=RAB-G3f | | | 0.012 | 0.018 | 0.028 | | | | | | |
| NET4B | tr\|Q84VY2\|Q84VY2_ARATH | At2g30500 OS=Arabidopsis thaliana GN=At2g30500 PE=2 SV=1 NET4B | 1.00 | 0.00 | 0.00 | 0.00 | 517 | 60287 | None | 10 | 7 | 16 |
| SD | | | 0.30 | 0.00 | 0.00 | 0.00 | | | | | | |
| P(plgem) | NH=RAB-G3f | | | 0.005 | 0.018 | 0.028 | | | | | | |
| VPS35B | sp\|F4I0P8\|VP35B_ARATH | Vacuolar protein sorting-associated protein 35B OS=Arabidopsis thaliana GN=VPS35B PE=1 SV=1 | 69.12 | 0.35 | 0.00 | 0.06 | 790 | 89514 | None | 326 | 56 | 77 |
| SD | | | 12.42 | 0.61 | 0.00 | 0.12 | | | | | | |

| | | | | | | | | | | | | |
|---|---|---|---|---|---|---|---|---|---|---|---|---|
| P(plgem) | NH=RAB-G3f | | | 2.0E-05 | 1.9E-05 | 4.70E-05 | | | | | | |
| TRS120 | tr\|Q9FY61\|Q9FY61_ARATH | Protein TRS120 OS=Arabidopsis thaliana GN=T5K6_30 PE=4 SV=1 | 0.56 | 7.81 | 6.47 | 0.31 | 1186 | 129712 | None | 281 | 62 | 61 |
| SD | | | 0.38 | 7.22 | 5.08 | 0.31 | | | | | | |
| P(plgem) | NH=RAB-G3f | | | 1.58 E-03 | 3.44E -03 | 0.47 | | | | | | |
| | NH=RAB-A2a | | 1.58E-03 | | 0.34 | 3.57E-04 | | | | | | |
| | | | n = 3 | n = 5 | n = 3 | n = 6 | | | | | | |

*SINQ quantification values all divided by 0.0000002759

**Table S3** Potential RABG3b interactors retrieved from yeast-two-hybrid screening.

| **ATG Code** | **Description** | **No. Clones** |
|---|---|---|
| AT5G58320 | AtNET4A | 20 |
| AT3G19900 | Uncharacterised Protein | 19 |
| AT1G48540 | Outer arm dynein light chain 1 protein | 15 |
| AT5G41790 | CIP1 (COP1-INTERACTIVE PROTEIN 1) | 7 |
| AT2G25730 | Uncharacterised Protein | 6 |
| AT2G14680 | MATERNAL EFFECT EMBRYO ARREST 13 | 6 |

**Table S4. Mutants used in this study**

| ATG number | Gene name | T-DNA line |
|---|---|---|
| At4g09720 | RABG3a | SALK_139519C |
| At2g22740 | RABG3b | SALK_004938 |
| At3g16100 | RABG3c | SALK_021190C |
| At1g52280 | RABG3d | GABI_967G07 |
| At1g49300 | RABG3e | SALK_058079C |
| At3g18820 | RABG3f | SALK_122061C |
| At2G21880 | RABG2 | SALK_069603C |
| AT5G58320 | NET4A | SAIL_116_C08 |
| | | SALK_083604 |
| At2g30500 | NET4B | SALK_056957 |

**Table S5.** PCR primers used in this study for gene expression analysis

| Gene name | ATG number | Forward primer 5'->3' | Reverse primer 5'->3' |
|---|---|---|---|
| U-Box | AT5G15400 | TGCGCTGCCAGATAATACACTATT | TGCTGCCCAACATCAGGTT |
| RABG3A | At4g09720 | CTTGATGAATCAATATGTGCATAA | CATCATAAACCAAAGCACAAC |
| RABG3B | AT1G22740 | AATAGCCGAGTGGTATCTGAGA | AACCAGTATCTGGCTGGAAATAT |
| RABG3C | At3g16100 | GAGTCGAGTGGTTACTGAGAA | ACTTCTTCTTCAGGTTCATTCT |
| RABG3D | At1g52280 | TCTCAGAAACCACTCTGCTCT | GCTGGACAAGAAAGATTTCAA |
| RABG3E | At1g49300 | ACTTTACAGATCTGGGACACA | GAAAGTTCTCTGGATCCGAAG |
| RABG3F | At3g18820 | CTTATCGGAAACAAGGTTGAC | GAAAGCTTCCTCCACATTAGT |
| NET4A | AT5G58320 | ATTATGATCTGCTTCGTTCCA | CTCCTCTATCAACTTCACCATC |
| NET4B | At2g30500 | TAGCTTGATCATAACGCTCAG | AATCTAGAAAAGATGGATGATCG |
| net1b.1 | At2g30500 | CGGTGACGAGGCATTGATCCG | CTCGTCTTCGTGCATTCGCA |
| net1a.1 | AT5G58320 | ATGGATTATGATCTGCTTCGTTCCAAG AAG | GGTGGATGCCCTGAAAAGGC |
| EF1a Fw | AT2G18720 | CCCATTTGTGCCAATCTCT | CACCGTTCCAATACCACCAA |
| TIP41 | At4g34270 | GTGAAAACTGTTGGAGAGAAGCAA | TCAACTGGATACCCTTTCGCA |

**Supplemental videos**

**Video/Movie S1** Confocal microscopy revealed discrete NET4A-GFP punctae labelling along the Lifeact-RFP actin filaments. Movie shows fly through of merged channel z series.

**Video/Movie S2** Arabidopsis lines co-expressing native promotor-driven NET4A-GFP and RABG3f-mCherry showed that NET4A localizes at the tonoplast  Movie shows fly through of merged channel z series.

**Video/Movie S3** Arabidopsis lines co-expressing native promotor-driven NET4A-GFP and RABG3f-mCherry showed that NET4A localizes at the tonoplast. Movie shows a maximum intensity 3D render of a z series focusing on two vacuoles. First the NET4A-GFP signal is shown, then the RABG3f-mCherry signal, then a merge of the two channels.
